# Pupil Dilation and Response Slowing Distinguish Deliberate Explorative Choices in the Probabilistic Learning Task

**DOI:** 10.1101/2021.10.19.464963

**Authors:** Galina L. Kozunova, Ksenia E. Sayfulina, Andrey O. Prokofyev, Vladimir A. Medvedev, Anna M. Rytikova, Tatiana A. Stroganova, Boris V. Chernyshev

**Author notes:** Data are openly available at https://doi.org/10.6084/m9.figshare.16825570. Correspondence concerning this article should be addressed to Boris V. Chernyshev, Center for Neurocognitive Research (MEG-Center), Moscow State University of Psychology & Education, 29 Sretenka str., Moscow, 127051, Russia.

## Abstract

This study examined whether pupil size and response time would distinguish directed exploration from random exploration and exploitation. Eighty-nine participants performed the two-choice probabilistic learning task while their pupil size and response time were continuously recorded. Using LMM analysis, we estimated differences in the pupil size and response time between the advantageous and disadvantageous choices as a function of learning success, i.e., whether or not a participant has learned the probabilistic contingency between choices and their outcomes. We proposed that before a true value of each choice became known to a decision-maker, both advantageous and disadvantageous choices represented a random exploration of the two options with an equally uncertain outcome, whereas the same choices after learning manifested exploitation and direct exploration strategies, respectively. We found that disadvantageous choices were associated with increases both in response time and pupil size, but only after the participants had learned the choice-reward contingencies. For the pupil size, this effect was strongly amplified for those disadvantageous choices that immediately followed gains as compared to losses in the preceding choice. Pupil size modulations were evident during the behavioral choice rather than during the pretrial baseline. These findings suggest that occasional disadvantageous choices, which violate the acquired internal utility model, represent directed exploration. This exploratory strategy shifts choice priorities in favor of information seeking and its autonomic and behavioral concomitants are mainly driven by the conflict between the behavioral plan of the intended exploratory choice and its strong alternative, which has already proven to be more rewarding.

---

“A bird in the hand is worth two in the bush” – will you follow this common wisdom, or will you ever abandon something good you already have and venture into the unknown in the vague hope of a bigger win? In a probabilistic environment, people usually tend to imagine hidden regularities in the outcomes of their actions – even when no such regularities actually exist (Ellerby & Tunney, 2017; Unturbe & Corominas, 2007). Attempting to test their surmises and catch a lucky break, people explore apparently disadvantageous options instead of just sticking to familiar profitable ones (Shanks, Tunney, & McCarthy, 2002). In fact, by doing so in probabilistic experimental tasks involving truly random and mutually independent choice outcomes, they actually fail to maximize their profits (Guttel & Harel, 2005; Unturbe & Corominas, 2007; Vulkan, 2000).

In contrast to typical artificial experimental conditions, outcomes of one’s actions in real life may be nonrandom and interdependent, e.g., the outcome of the next trial may be a consequence of the outcome of the previous trial. In this respect, exploring uncertain options instead of continuously exploiting a more rewarding alternative might bring new information about possible rewards, and thus might increase payoffs in the long run (Cogliati Dezza, Yu, Cleeremans, & Alexander, 2017; Sayfulina, Kozunova, Medvedev, Rytikova, & Chernyshev, 2020). From this perspective, the occasional switches from the prepotent value-driven advantageous response tendency to a disadvantageous choice in the probabilistic task may be considered as directed exploration – exploratory behavior that occurs when our desire for information overrides our need for reward. This crucial distinction between two qualitatively different types of exploration – directed and random exploration – has recently gained growing attention in literature (Payzan-LeNestour & Bossaerts, 2012; Schulz & Gershman, 2019; Schwartenbeck et al., 2019; Wilson, Geana, White, Ludvig, & Cohen, 2014; Zajkowski, Kossut, & Wilson, 2017). Directed exploration is intentional and specifically related to information-seeking targeted at the most uncertain option. On the contrary, random exploration is essentially a noisy response-generation process, leading to choices made by chance. Importantly, such type of exploration may be observed during early stages of reinforcement learning (Averbeck, 2015; Cogliati Dezza et al., 2017; Schulz & Gershman, 2019; Sutton & Barto, 1999), when all options are uncertain for the subject, or in conditions when the value of the most valuable option has decreased (Schwartenbeck et al., 2019) – thus also creating a need for learning the new rule of choice-reward contingency.

Random and directed forms of exploration are supposed to differ in underlying brain mechanisms (e.g., Warren et al., 2017; Zajkowski et al., 2017). Yet, the distinction between the two forms of exploration has not been accounted for in many previous physiological studies of exploration-exploitation dilemma.

For more than a decade, the adaptive gain theory (Aston-Jones & Cohen, 2005) has been an important theoretical background that guided many neurocognitive studies of the balance between exploitation and exploration. According to this theory, the locus coeruleus-noradrenergic (LC-NA) neuromodulatory system plays an essential role in regulating the balance between the two strategies (Aston-Jones & Cohen, 2005; Usher, Cohen, Servan-Schreiber, Rajkowski, & Aston-Jones, 1999). Specifically, this theory proposes that the tonic mode of the LC-NA neuromodulatory system promotes disengagement from the task and processing of task-irrelevant stimuli, thus creating a proper control state for exploring the new options. The increased neuromodulatory activity occurring over rather long timescale is accompanied by tonic pupil dilation, which is measured during pre-trial baseline, and it is believed to be a reliable proxy of LC-NA activation in the brain (Joshi & Gold, 2020). The tonic pupil dilation has been widely used to investigate the physiological basis of exploratory behavior (Gilzenrat, Nieuwenhuis, Jepma, & Cohen, 2010; Jepma, Beek, Wagenmakers, van Gerven, & Nieuwenhuis, 2010; Jepma & Nieuwenhuis, 2011).

In line with the concept of task disengagement central to the adaptive gain theory, many previous pupil studies of exploration-exploitation trade-off used behavioral tasks that explicitly forced participants to commit exploratory choices: typically that was achieved by decreasing the reward associated with the preferred choice (e.g., Daw, O’Doherty, Dayan, Seymour, & Dolan, 2006; Jepma & Nieuwenhuis, 2011; Payzan-LeNestour & Bossaerts, 2012). Such exploration related to task disengagement has important hallmarks of random exploration (Wilson, Bonawitz, Costa, & Ebitz, 2021) – e.g., as operationalized in Schwartenbeck et al. (2019). Yet little is known about the neurocognitive mechanisms associated with self-generated directed exploration targeted at information seeking (Zajkowski et al., 2017). In sharp contrast with random exploration, decision making during directed exploration focuses on active processing of choice–related information, thus emphasizing the deliberative process that requires the subject’s attention. Pupil size, as a component of phasic arousal, changes rapidly in response to cognitive operations underlying such attentional decision making (Poe et al., 2020). Specifically, recent research has shown that neural encoding of uncertainty or conflict can trigger fast task-evoked pupil dilation within the same trial in probabilistic reinforcement learning tasks (Van Slooten, Jahfari, Knapen, & Theeuwes, 2018), as well as in a target discrimination task (Gilzenrat et al., 2010). This suggests that phasic arousal reflected in phasic rather than tonic pupil changes may influence decision to deliberately choose an option with a more uncertain outcome.

In the current study, we aimed to investigate the observable physiological (pupil size) and behavioral (response time) concomitants of self-generated directed exploratory choices initiated by a participant on his or her own, i.e., in the absence of any external triggers and within a uniform series of trials under unchanging reward probabilities. As far as we know, the pupil-related arousal during directed exploration has never been addressed in the literature related to the exploration-exploitation dilemma.

We used a two-alternative repetitive choice task, with one option bringing more gains than losses, and the other one bringing more losses than gains. Participants were learning the task rules by trial and error. We primarily aimed to explore the distinction between the low-payoff (LP) and high-payoff (HP) choices, which were committed after a participant had learnt the choice-reward contingency. At this stage, participants had already acquired statistics of reward values associated with choice option – i.e., they possessed an internal utility model that guided them to prefer the HP option to the LP one. We reasoned that when choosing the option yielding a lesser (mathematical) expectation of the reward, people were engaged in an active effort to gather the information they are interested in (directed exploration), and they deliberately violated the acquired utility model.

In order to check this assumption, we contrasted LP choices made after successful learning to LP choices committed during the experimental blocks, in which participants failed to acquire a preference for the advantageous option, and thus did not possess any internal utility model relevant to the task rules.

Our basic predictions were related to the conflict/uncertainty that specifically characterizes the suboptimal LP choices violating the internal utility model. Indeed, there is a substantial evidence that in order to explore and gather information on a less certain, risky and potentially non-rewarding option, participants have to inhibit the tendency to choose a highly rewarded safe alternative (Cogliati Dezza et al., 2017; Daw et al., 2006). Assuming that the drive to commit the exploratory choice of a disadvantageous low-payoff (LP) option can be induced internally as a matter of actively probing the environment, we can predict, that the decision to commit a directed exploration would be accompanied by a conflict between seeking information concerning the uncertain options and seeking greater immediate profit associated with the other options as predicted by the internal utility model of the task.

On the basis of our hypothesis and previous pupil and RT studies of decision making (e.g., Cavanagh, Wiecki, Kochar, & Frank, 2014; Egner, 2007; Gilzenrat et al., 2010; Hershman & Henik, 2019; Laeng, Orbo, Holmlund, & Miozzo, 2011; Lin, Saunders, Hutcherson, & Inzlicht, 2018; Satterthwaite et al., 2007; Urai, Braun, & Donner, 2017; Van Slooten et al., 2018), we anticipated that conflict/uncertainty pertaining to self-generated exploratory choices would lead to greater phasic pupil-related arousal and RT slowing during the LP as compared to “safe” HP choices, but only when the decision-maker has learnt the choice-reward contingencies. We also expected that greater pupil dilation and decision costs would differentiate the LP choices made in the context of directed exploration from “random” LP choices committed under ‘no learning’ condition. Since tonic LC-NA activation measured via pre-trial pupil dilation has been implicated in exploratory behaviour (Gilzenrat et al., 2010; Jepma & Nieuwenhuis, 2011), we also investigated whether similar effects exist during self-generated directed exploratory behavior.

## Methods

### Participants

Ninety-four volunteers recruited from the community participated in the experiment (46 men and 48 women), aged 25.9 ± 5.6 years (*M* ± *SD*). All participants reported no neurological disorders or severe visual impairments; visual acuity was within ± 2.5 diopters, at least for a better-seeing eye. During the experiment, the participants did not use any vision correction devices (such as glasses or contact lenses).

The study was conducted following the ethical principles regarding human experimentation (Helsinki Declaration) and approved by the Ethics Committee of the Moscow State University of Psychology and Education. All participants signed the informed consent before the experiment.

### Procedure

During the experiment, participants were comfortably seated in an armchair and placed their heads on the chin rest to minimize involuntary head movements. We used the modified probabilistic learning task (Frank, Seeberger, & O’Reilly, 2004; Kozunova, Voronin, Venidiktov, & Stroganova, 2018), which was rendered as a computer game. On each trial, participants had to choose between two stimuli presented on the screen simultaneously. One stimulus was assigned as the advantageous (choosing this stimulus led to gains on 70% of trials and to losses on 30% of trials), and the other one as the disadvantageous (leading to gains on 30% of trials and to losses on 70% of trials). Probabilities were kept constant and did not change during the experiment. Losses and gains were assigned by a computer in a quasi-random order. Before the experiment, the participants were informed that the two stimuli were not equal in terms of the number of gains they could bring, yet no further information was revealed to the participants. Thus, the instruction could not prompt any specific choice strategies, and the participants had to learn from their own experience in trial-and-error fashion.

Each pair of stimuli comprised two images of the same Hiragana hieroglyph rotated at two different angles and rendered in white on black background (Figure 1). The size of the stimuli was 1.54 × 1.44°, which well fits into the fovea area. The stimuli were equalized in size, brightness, perceptual complexity, and spatial position. The two stimuli were located symmetrically on the left and on the right of the screen at 1.5° to the left and to the right from the screen center. The location of stimuli was alternated pseudorandomly from trial to trial with equal probability.

**Figure 1.**
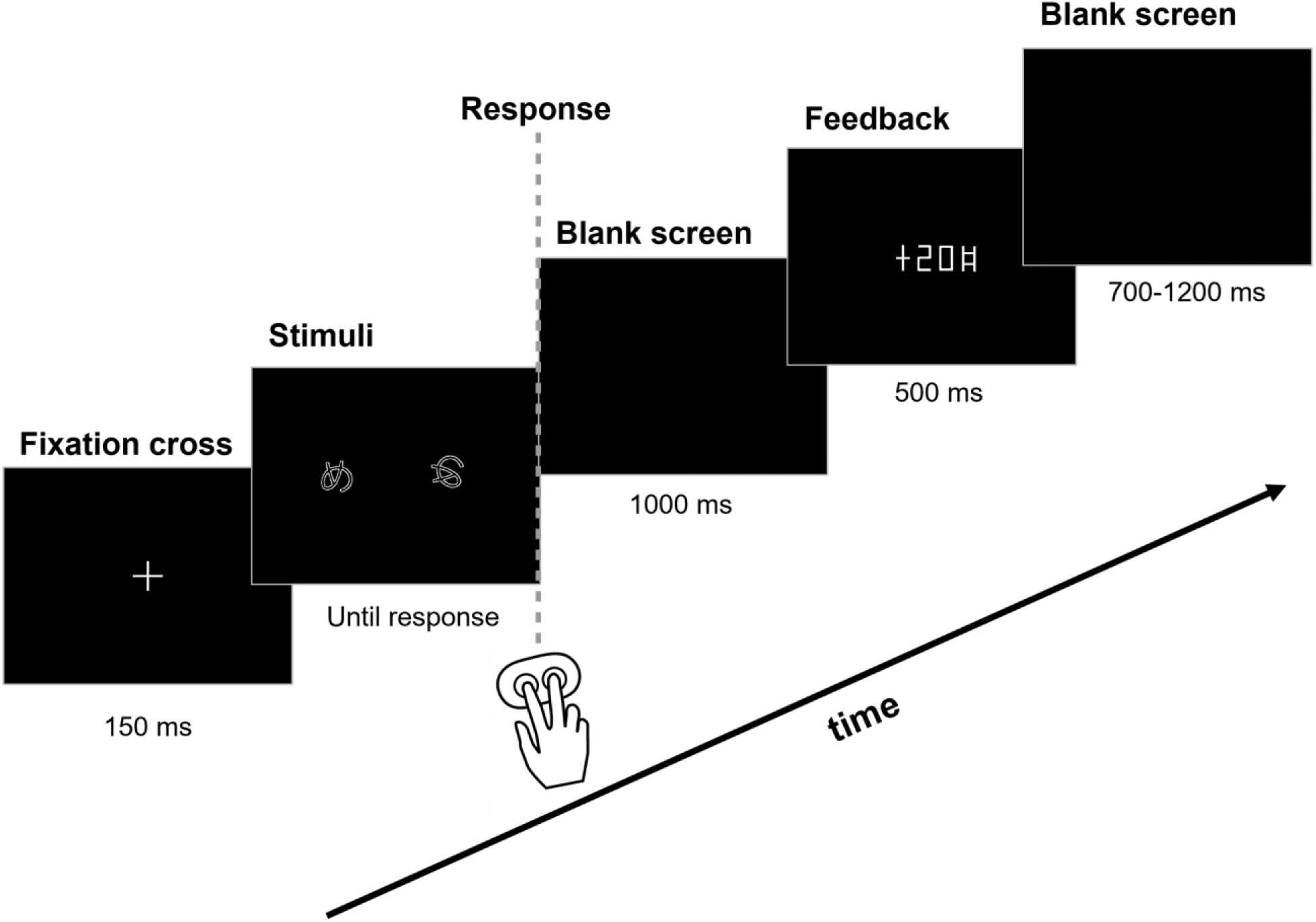
Probabilistic Value-Based Decision-Making Task. Each stimulus pair (a Hiragana hieroglyph rotated at two different angles) was presented repeatedly, and the left-right position of the stimulus on the screen was counterbalanced throughout an experimental block. Stimulus pairs and the expected value of each choice in a pair varied between the five blocks. Participants learned to select the more profitable of the two options (the advantageous one) solely from probabilistic feedback. After each trial, the number of points earned was displayed after a 1000 ms delay after a participant made a choice of one of the options by pressing the left or right button (for the left and right choices respectively). See text for details.

Before the start of the experiment, participants completed a quick test for visual discrimination and recognition of figures similar to those used during the experiment. All participants passed the test well.

On each trial, before the stimulus onset, a white fixation cross on black background was presented for 150 ms (Figure 1). Stimuli remained on the screen until a button was pressed by the participant; the instruction given to the participants did not require keeping gaze fixation during stimulus presentation, and the participants were allowed to freely view the stimuli. Aiming to avoid time pressure on participants and thus minimize the number of impulsive decisions, we did not impose any time limit on the response time.

During the experiments, participants continuously kept their index and middle fingers of the right hand on two buttons of the gamepad. To choose one of the two stimuli, participants pressed one of the buttons according to the location of the chosen stimulus on the screen (i.e., they pressed the left button to choose the left stimulus and the right button to choose the right one).

Immediately after the button press, the screen was cleared and remained empty (black). After a delay of one second following the button press, the visual feedback was presented for 500 ms. The feedback informed the participants about the number of points they received or lost on the current trial (Figure 1). These points were accumulated throughout the experiment; at the end of each block, participants were shown their total accumulated score. The screen was black during the intertrial interval.

We used an intertrial interval ranging from 700 to 1400 ms, which varied in a quasi-random order (flat distribution). We used a rather short intertrial interval, thus keeping the duration of the whole experiment to a minimum. This was done with the aim of preventing fatigue and boredom in participants and maintaining their interest and motivation. This intertrial interval was shorter than the duration of the pupillometric effects; we addressed this issue at the stage of preprocessing the pupillometric data (see below). The intertrial interval was varied in order to prevent rhythmical responding that could lead to impulsive or perseverative responses.

The experiment involved five similar blocks. Each block included 40 trials and lasted about 5 minutes. A short rest lasting about 1 minute was introduced between blocks. The total duration of the experiment was about 30 minutes. A new stimulus pair was used in each block. Although the probabilities of gains and losses associated with the advantageous and disadvantageous stimuli were kept constant throughout the whole experiment, blocks differed in the exact number of points assigned as gains and losses. We used five reinforcement schemes with the following magnitudes of gains and losses, respectively: (I) +20 & 0, (II) 0 & −20, (III) +50 & +20, (IV) −20 & −50, (V) +20 & −20. Changing reinforcement scale between blocks was needed in order to make the blocks appear less similar and thus to make the participants learn during each block anew.

The order of reinforcement schemes within the blocks was counterbalanced across participants by means of using three different sequences: I–II–III–IV–V, V–III–II–IV–I and V– III–II–I–IV. The sequences were assigned to participants randomly. We did not use other possible combinatory variants of sequences in order to keep the accumulated score in all participants above zero throughout the whole experiment duration. Maintaining a positive balance was needed to prevent participants from experiencing frustration and losing motivation. The accumulated score was converted from points to rubles at 1:1 ratio, and the money was paid to participants after the experiment. On average, participants were paid 420 ± 250 rubles (*M* ± *SD*).

The experiment was implemented using the Presentation 14.4 software (Neurobehavioral systems, Inc., Albany, CA, USA).

### Pupillometric Recording

Pupil size was recorded continuously from the participants’ dominant eye using an EyeLink 1000 Plus infrared eye tracker (SR Research Ltd., Canada) at the sampling rate of 1000 Hz. Pupil size was measured as pupil area in camera pixels using the default eye tracker settings. Before each block, participants completed the EyeLink 1000 9-point calibration procedure.

### Data Preprocessing

Response time (RT) was measured as an interval from stimulus onset to a button press. Trials with extreme RT values (<300 ms and >4000 ms) were excluded from the analysis; such trials comprised 4.7% of the experimental data. After that, RT data from all valid trials in all blocks jointly were z-transformed within each subject in order to reduce inter-subject variability in RT and make the distribution closer to normal.

Preprocessing of pupillometric data was performed with custom-made scripts in R Studio (Version 3.6.3; R Core Team, 2020) using the ‘eyelinker’ package (Barthelme, 2019). Artifacts related to short interruptions or malfunctioning of pupillometric recording due to blinks or other causes were detected using the following criteria: pupillometric data were missing or the absolute value of the derivative of the pupillometric data exceeded 10 pixels between two adjacent measurements (which were recorded at 1 ms steps). For relatively short artifacts up to 350 ms, which were caused by ordinary eyeblinks, data were replaced by linear data interpolation. If the artifact duration exceeded 350 ms, the respective 5-second epochs (−2000 – 3000 ms relative to the button press) were excluded from the analysis of the pupillometric data.

Next, we downsampled pupillometric data to 20 time points per second by averaging the data into consecutive 50-ms intervals.

After that, in order to reduce inter-subject variability in pupil size and make the distribution closer to normal, pupil data were z-transformed within each subject. For this purpose, we used continuous pupillometric data from all five experimental blocks jointly. After that, we excluded from the analysis those epochs during which z-transformed pupil size deviated from zero by more than 3 standard deviations; such trials comprised 0.56% of the experimental data. All pupillometric measurements are reported below as z-scores.

After z-scoring, the relative zero of pupillometric measurements represented an average throughout all trials of the experiment in each participant, one and the same for all conditions and timepoints. We used this normalization procedure because the intertrial intervals in the current experiment were rather short compared with the duration of “cognitive” pupillometric responses, which can last up to several seconds (e.g., de Gee, Knapen, & Donner, 2014; Koenig, Uengoer, & Lachnit, 2018; Lavin, San Martin, & Jubal, 2014; Preuschoff, Hart, & Einhauser, 2011). Z-scores allowed us to account for carryover effects between adjacent trials, while a conventional baseline correction by means of subtracting the “pretrial baseline” could lead to erroneous estimation of phasic pupil responses. Particularly, one cannot exclude that baseline-corrected pupil size can be influenced by the long-lasting pupil changes occurring on a preceding trial and lasting throughout the intertrial interval. A common measurement scale for all conditions allowed us to make direct comparisons between conditions and timepoints, with a higher z-score value always reflecting greater pupil size. Note, that the trial-related z-scores might be small or negative because the highest positive z-scores were always obtained during the pre-trial baseline characterized by pupil dilation in the absence of any visual stimulation on the screen. Additionally, in order to evaluate slow non-phasic effects, we analyzed the pretrial pupil size (see below). We also supplemented this report by repeating the main analyses using the pupil size data that were baseline-corrected by means of subtracting the pretrial pupil size (see below).

### Trial Selection Criteria and the Factors of Interest

We assumed that when participants exhibited a stable preference for the advantageous stimulus, they guided their behavior using the utility model that they had acquired through trial-and-error learning during the initial trials of a given block. Therefore, after learning, disadvantageous choices were likely committed against the internal utility model, and thus they could represent intentional directed exploration. Within each block independently, we first identified all trials belonging to such ‘after learning’ periods using the following criteria:

1. Such periods should be preceded by four advantageous choices made in an interrupted succession, and
2. The percentage of advantageous choices thereafter until the end of the block should be no less than 65%. We will refer hereafter to such periods as ‘after learning’ condition.

Then, we identified trials during which the participants made objectively advantageous choices and disadvantageous choices (i.e., when they selected the stimuli associated with 70% and 30% gain probability, respectively).

Potentially, the disadvantageous choices could be caused solely by negative outcomes of a preceding advantageous choice, which could prompt the participants to immediately change their preference (Gaffan & Davies, 1981; Ivan, Banks, Goodfellow, & Gruber, 2018) – i.e., to follow a Win-Stay Lose-Shift strategy (see e.g., Ellerby & Tunney, 2017). In order to check whether the disadvantageous choices were mainly caused by a previous loss, we investigated probabilities of transitions leading from the advantageous to the disadvantageous choice. We took the number of all transitions from advantageous stimuli to disadvantageous ones as 100% and calculated the percentage of transitions made after losses. Then we used a t-test to compare this percentage to 100% (as if all transitions to exploration were made strictly as a response to losses) and to 30% (as if transitions to exploration were strictly independent of the preceding feedback).

Next, in order to investigate the dynamics of behavioral and pupillometric indices over transitions between types of choices made by participants, we used a more detailed trial classification procedure. For this purpose, we took into account not only the choice made during each given trial but also the choices made on the previous one and on the following one. Not all possible combinations could be encountered often enough: only four combinations had a sufficient number of trials during ‘after learning’ condition (>6 trials for each condition per subject on average), while other possible combinations were rather rare (<2 trials for each condition per subject on average). Thus, for further analyses, we used the following four levels of ‘Choice Type’ factor (Table 1):

- the ‘low-payoff’ choice (LP) – the disadvantageous choice preceded and followed by advantageous choices;
- the ‘high-payoff’ choice (HP) – the advantageous choice preceded and followed by advantageous choices, thus representing a stable preference for advantageous stimuli;
- the trial preceding the ‘low-payoff’ choice (pre-LP) – the advantageous choice that preceded the disadvantageous and followed the advantageous one;
- the trial following the ‘low-payoff’ choice (post-LP) – the advantageous choice that followed the disadvantageous and preceded the advantageous one.

**Table 1.** Choice Types and Overall Behavioral Statistics.

| Choice Type | Previous Trial →<br>Current Trial<br>→ Next Trial* | Number of Trials per Subject<br>($M \pm SD$) | | % of Trials per Subject<br>($M \pm SD$) | | Total Number of Trials | | RT (ms) | |
| --- | --- | --- | --- | --- | --- | --- | --- | --- | --- |
|  |  | After Learning | No Learning | After Learning | No Learning | After Learning | No Learning | After Learning | No Learning |
| HP | A → **A** → A | 55.8 ± 43.3 | 8.4 ± 7.0 | 60.1 ± 23.8 | 15 ± 9.7 | 4965 | 412 | 1171 ± 403 | 1577 ± 717 |
| pre-LP | A → **A** → DA | 7.7 ± 6.0 | 8.5 ± 6.2 | 11.8 ± 11.7 | 14.3 ± 6.2 | 686 | 416 | 1393 ± 606 | 1467 ± 597 |
| LP | A → **DA** → A | 6.6 ± 5.3 | 6.7 ± 5.8 | 9.7 ± 8 | 11.1 ± 7.0 | 587 | 330 | 1609 ± 597 | 1631 ± 703 |
| post-LP | DA → **A** → A | 7.2 ± 5.9 | 8.3 ± 5.6 | 9.8 ± 6.6 | 14.9 ± 5.7 | 639 | 406 | 1467 ± 610 | 1666 ± 669 |

\* A – advantageous choice, DA – disadvantageous choice.

As mentioned above, one could expect that the disadvantageous choices could be provoked by negative outcomes of the preceding advantageous choices, thus obscuring the supposedly exploratory nature of disadvantageous choices. In order to check whether this potentially retroactive mechanism did not influence the results substantially, we supplemented this report by repeating the basic analyses related to ‘Choice Type’ factor (see below and supplementary materials) on a smaller subset of trials with two additional restrictions:

1. The outcome of the pre-LP trial within such a sequence of trials should be a gain rather than a loss, and
2. Pre-LP, LP and post-LP trials should constitute uninterrupted sequences of trials in direct succession.

Most importantly for testing the validity of the “directed exploration” hypothesis, we contrasted the same types of choices performed during ‘after learning’ condition vs. choices made in ‘no learning’ condition (Learning factor). In the latter case, the respective trials were selected from blocks during which the participants failed to reach the learning criteria. We suggested that during such blocks, participants did not acquire a proper internal utility model, and any choices made by a participant, regardless its objective advantageousness, represented random rather than directed exploration.

### Statistical Analysis of RT and Pupil Size Using the Linear Mixed Effects Model

We used linear mixed effects models (LMM) at single-trial level rather than repeated measures ANOVA at the grand-average level because LMM method is robust to imbalanced designs across individual cases. Thus, missing data need not result in listwise deletion of cases, and differing numbers of trials per condition are less problematic than in traditional ANOVA (e.g., Kliegl, Wei, Dambacher, Yan, & Zhou, 2011). LMM can handle large numbers of repeated measurements per participant, thus making it possible to analyze data from individual trials: this allows accounting for intertrial variability, which would be lost under standard averaging approaches (Tibon & Levy, 2015; Vossen, Van Breukelen, Hermens, Van Os, & Lousberg, 2011).

Statistical LMM analyses were performed using R software v 4.1.0 (R Core Team, 2021)

We used the following basic procedure, unless specified otherwise. We fitted LMMs on RT and pupillometric data using lme4 package (Bates, Mächler, Bolker, & Walker, 2015). We started with the full model, which included relevant fixed effects and their interactions. In the current report, we used the following fixed effects and their interactions: ‘Choice Type’ (four levels, ‘HP’, ‘pre-LP’, ‘LP’, and ‘post-LP’) – as described above, ‘Previous Feedback’ (two levels, ‘gain’, and ‘loss’ – the outcome of the trial that immediately preceded the current one), ‘Current Feedback’ (two levels, ‘gain’, and ‘loss’ – the outcome of the choice made during the current trial), and ‘Learning’ (two levels, ‘after learning’, and ‘no learning’ – as described above).

All models used for data analysis in the current study included the following random effects intercepts: ‘Subject’ (89 levels), ‘Block number’ (five levels, ‘one’ to ‘five’ – position of a particular block in the sequence of experimental blocks), and ‘Reinforcement Scheme’ (five levels, the monetary value of gains and losses within a particular experimental block, see above).

The LMMs for the main analyses included the following fixed factors: ‘Choice Type’, ‘Previous Feedback’ and ‘Learning’:

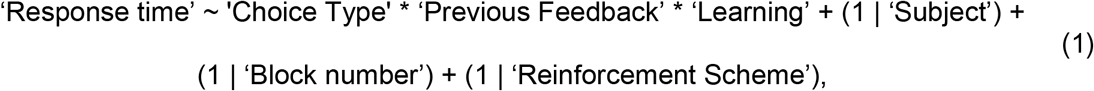

and

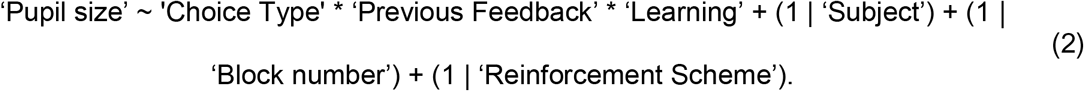

Additionally, for illustrative purposes, we aimed to look into the ‘Choice Type’ x ‘Previous Feedback’ interaction, within ‘no learning’ and ‘after learning’ conditions analyzed separately. For this purpose, we used the following LMMs for RT and pupil size:

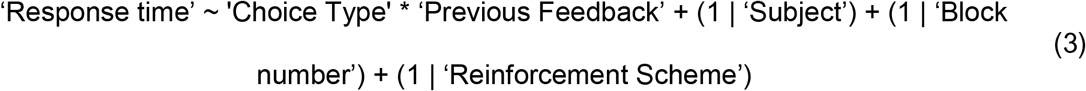

and

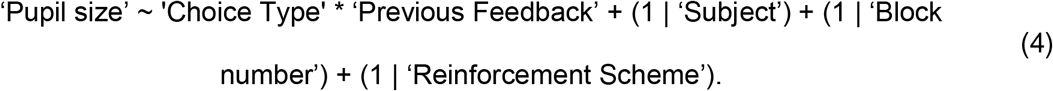

In the same vein, when ‘Learning’ factor did not interact with the ‘Choice Type’ and ‘Current Feedback’, we illustrated the ‘Choice Type’ x ‘Current Feedback’ interaction, within ‘no learning’ and ‘after learning’ conditions analyzed separately. For this purpose, we used the following LMM for the analysis of the pupil size when focusing on the time interval after the feedback onset:

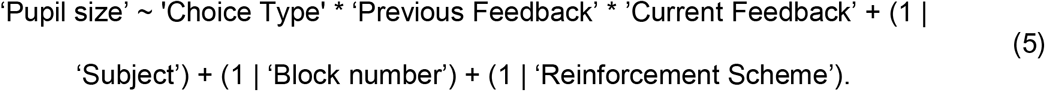

For all models, we did a step-down model selection procedure using ‘step’ function implemented in lmerTest package (Kuznetsova, Brockhoff, & Christensen, 2017). This procedure performs backward elimination of nonsignificant effects. At the first stage, nonsignificant random effects are eliminated based on of the likelihood ratio test (Stuart, Ord, & Arnold, 1999). Then significance of fixed factors is assessed using the Kenward-Roger approximation for denominator degrees of freedom (Halekoh & Højsgaard, 2014; Kuznetsova et al., 2017). At each step, the nonsignificant factor with the highest *p*-value is eliminated, and this procedure is repeated until only significant factors remain in the model. Next, we used the simplified model produced by the ‘step’ function. For those factors that remained in the model, we estimated significance using the Satterthwaite approximation for denominator degrees of freedom and obtained type III ANOVA table (package lmerTest, function anova) (Kuznetsova et al., 2017).

Next, in order to investigate particular contrasts, we performed planned comparisons using the Tukey HSD post-hoc tests (Tukey, 1977) implemented in ‘emmeans’ package (Lenth, 2021). We did that in two ways. First, we evaluated pairwise differences within the levels of ‘Choice Type’ factor. Second, in order to investigate the interference between ‘Choice Type’ and other factors of interest (‘Previous Feedback’, ‘Current Feedback’, or ‘Learning’), we evaluated pairwise differences within the levels of the factor of interest split by the levels of ‘Choice Type’ factor.

### Defining the Time Interval of Interest for Testing the Effects of Learning and Previous Feedback on Pupil Size

First, we wanted to check whether the pupil size differentiated between advantageous and disadvantageous choices during ‘after learning’ condition – and, if it did, to determine the time span of these choice-specific pupillary responses. The average pupil size measured across the time interval selected this way would allow for testing the role of learning and/or previous feedback in the choice-driven pupil size modulations.

First, we analyzed independently each time point within the epoch ranging from − 1000 to 2500 ms relative to time of the response. We ran the following LMM on single-trial data with ‘Choice Type’ factor (four levels: HP, pre-LP, LP, post-LP) taken as the fixed effect:

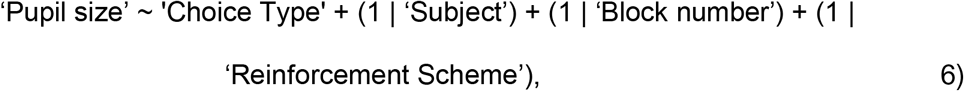

Next, we performed planned comparisons using the Tukey HSD post-hoc test within ‘Choice Type’ factor (four levels: HP, pre-LP, LP, post-LP), thus obtaining the statistical significance of pairwise differences between ‘Choice Type’ factor levels and applied correction for multiple comparisons using the false discovery rate (FDR) method for 70 time intervals at q=.05.

As a result, we obtained the time spans of significant differences in pupil size between choice types. Since we were interested in comparing the disadvantageous choices and neighboring trials vs. advantageous trials, we restricted this analysis to three contrasts: pre-LP choices vs. HP choices, LP choices vs. HP choices and post-LP choices vs. HP choices. For the use in further analyses, we chose the overlap of significant time intervals within these three contrasts. This procedure produced a rather long time interval from - 400 ms to 2200 ms relative to the behavioral response. Thus, for further analyses, pupil size was averaged within this interval, in each trial separately.

Apparently, this rather long interval was functionally heterogeneous. Thus, additionally, we divided this time interval into three functionally different subintervals, and analyzed them separately (see supplementary materials):

- decision making and action initiation (−400 – 0 ms relative to the response),
- internal outcome evaluation and feedback anticipation (0 – 1000 ms relative to the response),
- matching expected and actual feedback (1000 – 2200 ms relative to the response).

In order to evaluate slow non-phasic changes in pupil-size, that might partly reflect tonic effects, we also analyzed pretrial pupil size, averaged over −300 – 0 ms relative to the fixation cross onset. We also repeated the main analyses using the conventionally baseline-corrected data on a trial-to-trial basis; this correction was applied to z-transformed data by means of subtracting pretrial pupil size.

### Correlational Analysis

In order to test whether commission of disadvantageous choices decreases the profit gained by participants, we calculated Pearson’s correlation between the percentage of LP choices and the number of gains, both measures evaluated within ‘after learning’ condition. The percentage of disadvantageous choices was calculated for each participant using the following formula:

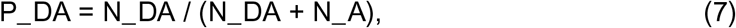

where N_DA is the number of all disadvantageous choices, and N_A is the number of advantageous choices for each participant within ‘after learning’ condition.

In addition, we aimed to investigate whether the effects related to disadvantageous choices under ‘after learning’ condition would be dependent upon how often the participants ventured into such disadvantageous choices. For this purpose, we investigated the relation between the percentage of disadvantageous choices and LP-choice-related changes in both RT and pupil size using Pearson’s correlation. LP-choice-related RT slowing was calculated for every subject as the difference between mean RT during LP choices and mean RT during HP choices during ‘after learning’ condition. LP-choice-related pupil dilation was calculated for every subject as the difference between mean pupil size during LP choices and mean pupil size during HP choices within a post-feedback time interval (1000 – 2200 ms relative to the behavioral response) during ‘after learning’ condition. In order to reduce statistical noise, we included onto this analysis only those 80 participants who had more than one LP choice.

We plotted scatterplots and respective linear regressions in order to illustrate the results of the correlational analyses. All statistical analyses were performed using R software v 4.1.0 (R Core Team, 2021).

## Results

### General Behavioral Statistics

Before the start of the experiment, participants successfully completed the test for discrimination within pairs of stimuli similar to those used in the experiment. This excludes the possibility that participants could have had any substantial difficulty in perceptual discrimination between the stimuli during the experiment. The overall behavioral statistics is shown in Table 1.

Over the entire period of the experiment (five blocks), the participants made disadvantageous LP choices on 27.1% ± 14.2% of trials (*M* ± *SD*). Eighty-nine out of 94 participants fulfilled the learning criteria (four advantageous HP choices committed consecutively and no less than 65% of advantageous HP choices thereafter until the end of the block) in at least one or greater number of experimental blocks. Since five participants completely failed to learn in any of the experimental blocks, they were excluded from all further analyses. The remaining 89 participants reached learning criteria on 3.7 ± 1.4 blocks (*M* ± *SD*) out of five. Learning criteria were reached by them after 12.4 ± 6.8 trials (*M* ± *SD*) out of 40 trials comprising each block.

Participants made disadvantageous choices on 17.0% ± 9.5% of trials within ‘after learning’ condition, and significantly more often (24.3% ± 22.2%) within ‘no learning’ condition (*t*_(88)_=3.00, *p*=.004).

When the stimulus-reward contingency was learned, 42.2% ± 25.3% of all transitions from advantageous to disadvantageous choices were committed after losses. This value is significantly smaller than 100% - the percentage that would have been observed if disadvantageous LP choices were triggered exclusively by losses (*t*_(86)_=-21.28, *p*<.001), and significantly greater than 30% – the percentage of negative outcomes of advantageous choices in the experimental procedure (*t*_(86)_=4.50, *p*<.001). This means that a previous loss, even if it violated the acquired utility model, did not fully account for the following choice of the disadvantageous option.

Within ‘no learning’ condition, a very similar pattern was observed: 39.2% ± 19.9% of all transitions from advantageous to disadvantageous choices were committed after losses. The pattern of results was quite similar to that observed ‘after learning’: the percentage of transitions after losses was significantly smaller than 100% (*t*_(52)_=-22.2, *p*<.001) and significantly greater than 30% (*t*_(52)_=3.29, *p*<.001).

The total number of gains negatively correlated with the percentage of LP choices (*r*_(74)_=-.57, *p*<.001), thus demonstrating that commission of disadvantageous choices indeed prevented participants from maximizing their cumulative profit, while the optimal strategy for them would be to avoid disadvantageous choices as much as possible (Figure 2a).

**Figure 2.**
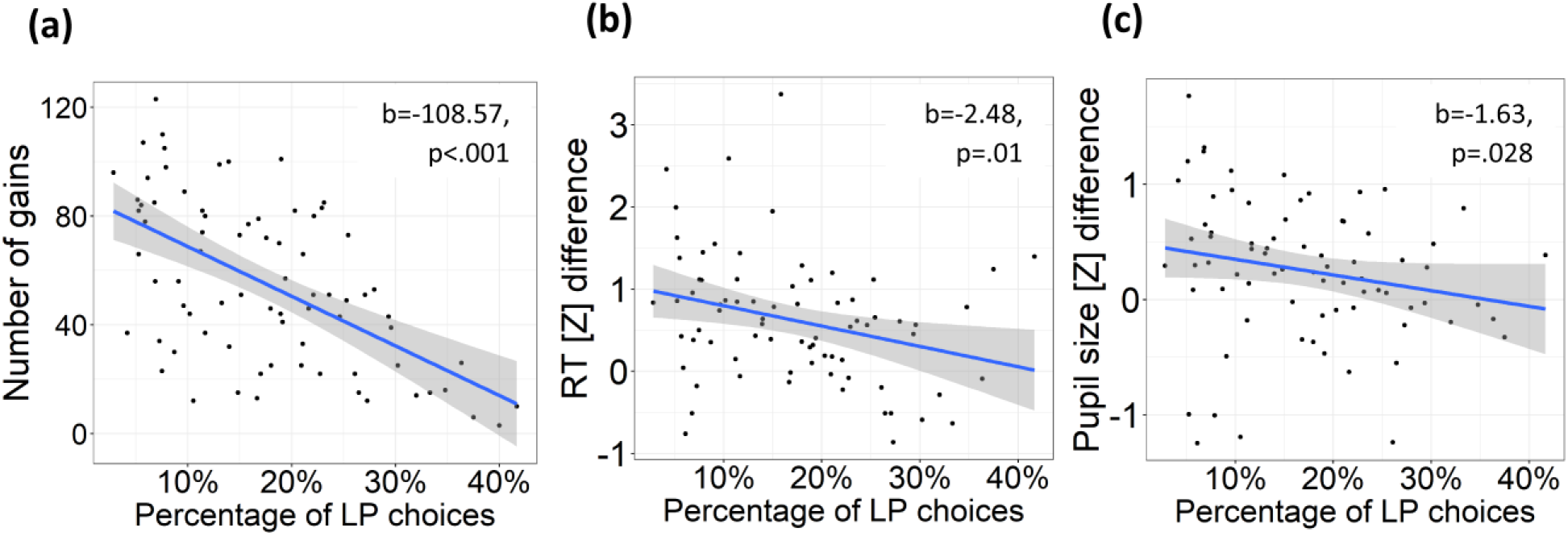
Scatterplots Depicting the Relationship Under ‘After Learning’ Condition. **(a)** between the percentage of LP choices and the number of gains obtained by participants; **(b)** between the percentage of LP choices and RT slowing during LP choices (difference in z-scores between LP and HP choices); **(c)** between the percentage of LP choices and relative pupil dilation during LP choices (difference in z-scores between LP and HP choices, averaged over 1000–2200 ms relative to response). Each dot on the scatterplots represents averaged data for one participant. Lines on scatterplots represent respective linear regressions, with shaded areas depicting 95% confidence intervals.

### Response Time: Effects of Learning and Previous Feedback

The LMM for the RT analysis included the following fixed effects: ‘Choice Type’ (HP, LP, pre-LP, and post-LP), ‘Previous Feedback’ (gains and losses), ‘Learning’ (‘no learning’ and ‘after learning’), and their interactions. The following effects and interactions were statistically significant: ‘Choice Type’ (*F*_(3,8187)_=26.2, *p*<.001), ‘Previous Feedback’ (*F*_(1,8106)_=6.6, *p*=.01), ‘Learning’ × ‘Choice Type’ (*F*_(3,8266)_=15.5, *p*<.001), ‘Choice Type’ × ‘Previous Feedback’ (*F*_(3,8340)_=5.99, *p*<.001).

#### Learning effect

Planned comparison revealed that ‘Learning’ × ‘Choice Type’ interaction was due to the fact that learning affected the RT for the LP and HP choices in opposite directions (Figure 3a, left panel): it slowed RT during LP choices (‘after learning’ vs ‘no learning’: Tukey HSD, *p*<.001) and accelerated RT during HP choices (‘after learning’ vs ‘no learning’: Tukey HSD, *p*<.001). Thus, learning of stimulus-reward contingency produced slowing of disadvantageous risky responses and speeding of advantageous HP responses that were committed during periods of stable preference for advantageous stimuli.

**Figure 3.**
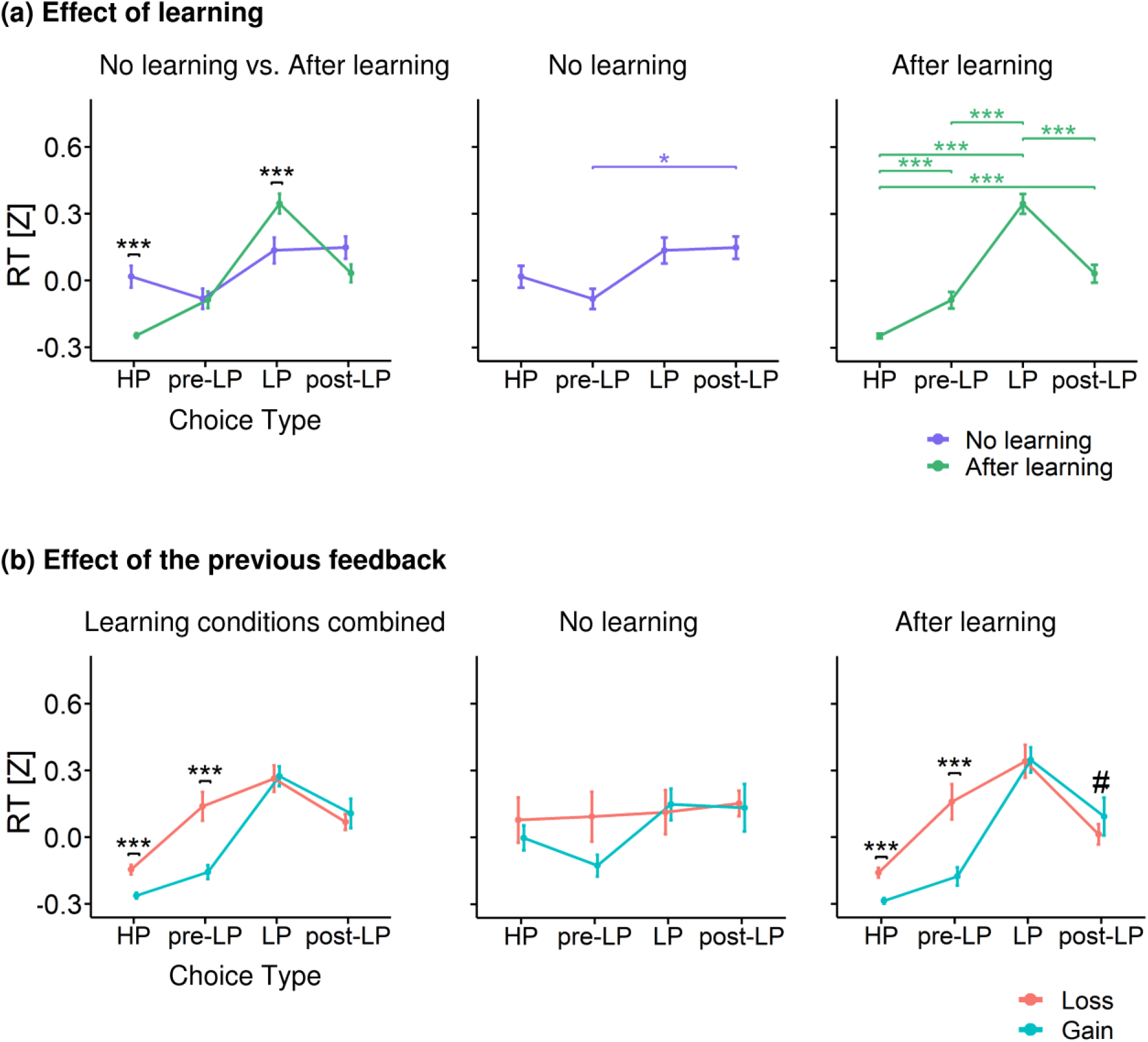
Response Time (z-scored) Represented as a Function of Choice Type. **(a)** RT for different choice types under ‘after learning’ (green) and ‘no learning’ (slate blue) conditions. Left panel – RT differences between learning conditions within each choice type, middle and right panels – RT differences between choice types within ‘no learning’ and ‘after learning’ respectively **(b)** RT differences between previous outcomes within each choice type: ‘losses’ (salmon) vs. ‘gains’ (turquoise). Left panel – both learning conditions pooled together, middle and right panels – ‘no learning’ and ‘after learning’ conditions respectively. Points and error bars on graphs represent M ± SEM across single trials in all subjects. # – p<0.1, * – p<0.05, ** – p<0.01, *** – p<0.001.

Next, we used planned comparisons within the same model to analyze ‘no learning’ and ‘after learning’ conditions separately. There were no significant differences in the RT between the LP and HP choice types in ‘no learning’ condition (Figure 3a, middle panel). On the other hand, after learning RT became significantly longer for the LP choices compared with all the other choice types, and significantly shorter for HP choices than RT for all the other choice types (Tukey HSD: *p*<.001 for pairwise contrasts between LP and HP, and for contrasts between HP/LP with pre-LP, and post-LP choices) (Figure 3a, right panel). Thus, in ‘after learning’ condition, but not in ‘no learning’ condition, decision making regarding disadvantageous choice took more time than that for all types of advantageous choices. Additionally, a moderate yet highly significant response slowing was observed on adjacent trials immediately preceding and immediately following a disadvantageous choice (pre-LP and post-LP choices).

In order to check whether the effect of RT slowing during disadvantageous LP choices was not a consequence of negative outcomes of a preceding advantageous choice, we repeated the same analysis using a smaller restricted subset of data within uninterrupted sequences of pre-LP → LP → post-LP trials involving only gains on pre-LP trials. The patterns of results concerning the effects of ‘Learning’ and ‘Choice Type’ were perfectly preserved in this reduced dataset (supplementary materials, Figure S1). Thus, the effects observed in relation to LP choices, did not result from losses on the preceding trial.

In summary, after learning, which led to formation of the internal utility model, two major changes occurred. First, RT decreased for stable preference for advantageous stimulus, revealing response speeding under a relatively safe strategy. Second, RT slowing was observed for a riskier strategy of disadvantageous choices.

#### Effect of previous feedback

When we considered ‘no learning’ and ‘after learning’ conditions together, we observed that losses and gains in the preceding trial differently affected RT for advantageous and disadvantageous choices (Figure 3b, left panel). RT was significantly slower after losses than after gains, but only in the case of advantageous HP and pre-LP choices (Tukey HSD: *p*’s <.001 for loss versus gain contrasts for both choice types). Thus, we observed post-loss slowing in situations, when both the previous choice and the current choice were advantageous; otherwise, there was no difference in RT between losses and gains.

Since the triple interaction ‘Choice Type’ × ‘Previous Feedback ‘× ‘Learning’ was not significant, we could not perform post hoc tests on the data split by all these factors. Instead, for illustrative purposes, we ran a similar LMM on ‘no learning’ and ‘after learning’ data subsets separately (Figure 3b, middle and right panels). ‘Choice Type’ × ‘Previous Feedback’ interaction was significant only within ‘after learning’ condition (*F*_(3,6846)_=4.74, *p*<.001). RT was significantly increased after losses compared with gains within HP choices and pre-LP choices (Tukey HSD: *p*’s <.001 for both comparisons). No significant differences were found within ‘no learning’ condition. Thus, the ‘Choice Type’ × ‘Previous Feedback’ interaction found in the full dataset was well pronounced mainly under ‘after learning’ condition, despite a similar trend under ‘no learning’ one.

### Pupil Size

#### Choice-driven Modulations of the Pupil Size Timecourses

We were primarily interested to find out whether the type of choice made by participants affected the pupil size. First, we needed to determine the time interval of interest for further analyses, and for this purpose, we ran the following analysis. At the first step, we analyzed each time point independently. We ran the LMMs on single-trial data with ‘Choice Type’ factor as a fixed effect. Next, we performed planned comparisons using the Tukey HSD post-hoc test for ‘Choice Type’ factor (four levels: HP, pre-LP, LP, post-LP). Since at the first step we analyzed each time point independently, at the second step we applied correction for multiple comparisons using the false discovery rate (FDR) method (Benjamini & Yekutieli, 2001) for 70 time points at q=.05.

The impact of factor Choice Type on pupil size timecourses under ‘after learning’ condition are represented in Figure 4. As contrasted with HP choices, the pupil was significantly larger during pre-LP choices (all qs <.05 (FDR corrected Tukey HSD) for the time interval from - 400 ms to 2200 ms relative to the button press, Figure 4, left panel), LP choices (all qs <.05 for the time interval from - 400 ms to 2200 ms, Figure 4, middle panel) and post-LP choices (all qs<.05 for the time interval from - 1000 ms to 2200 ms, Figure 4, right panel). Thus, for the use in further analyses, we took the pupil size averaged over the overlap of significant time intervals for all contrasts shown above, i.e., from - 400 ms to 2200 ms relative to the choice response.

**Figure 4.**
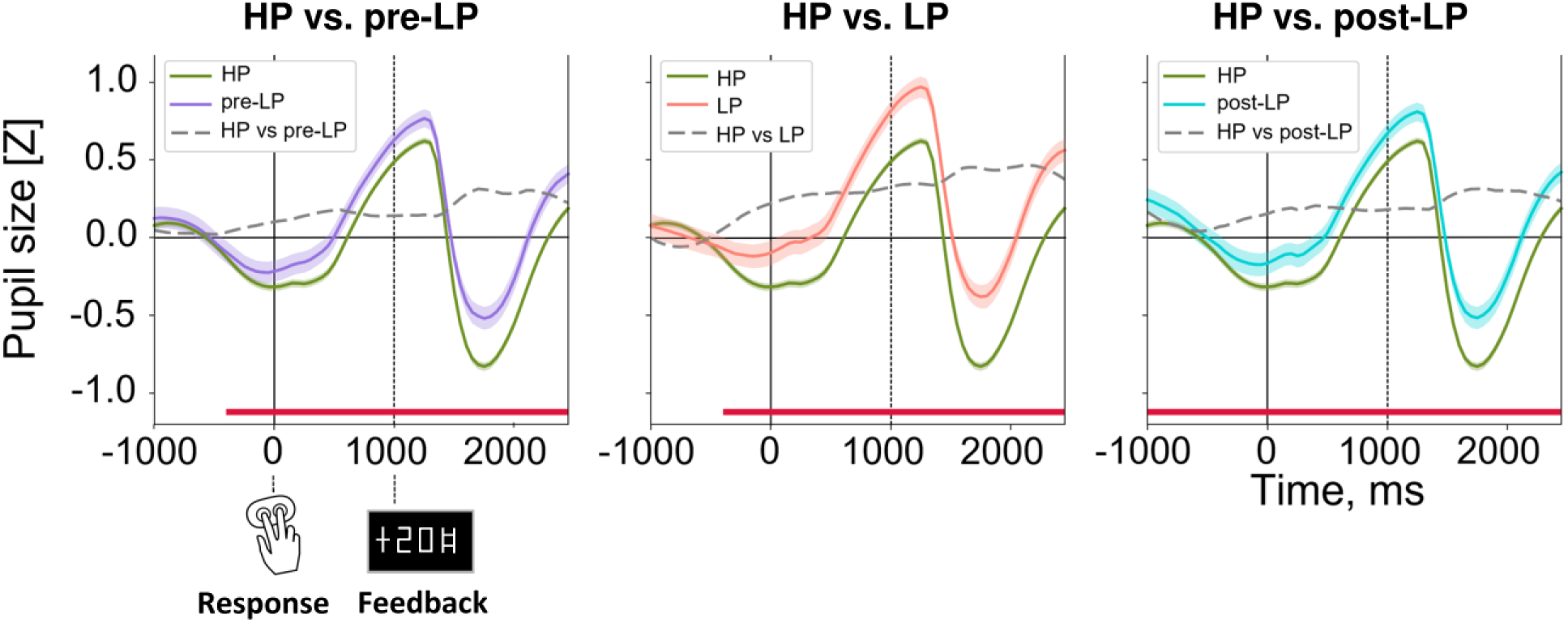
Time Courses of Pupil Size (z-scored) during Choice and after Feedback in the ‘After Learning’ Condition. From left to right: The pre-LP (violet), LP (red), and post-LP (marine blue) choices in comparison with the HP choice (green). The dashed curve in each graph corresponds to the time course of the difference in the pupil size between the respective choice type and the HP choice. Solid and dashed vertical lines correspond to button press (zero point) and feedback onset respectively. Curves and shaded areas represent *M* ± *SEM* across single trials in all subjects. Red lines at the bottom of each graph indicate significant differences between the respective choice type and HP choice (*p*<0.05, FDR corrected).

#### Effects of Learning and Previous Feedback

The model for this analysis included the following fixed effects: ‘Choice Type’, ‘Previous Feedback’, ‘Learning’ and interactions between them.

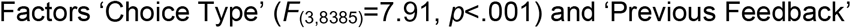

(*F*_(1,8422)_=16.27, *p*<.001) were significant (note, that significance for ‘Choice Type’ factor was partially related to the method we used to define the time interval for the analysis). More importantly, there were statistically significant interactions: ‘Learning’ × ‘Choice Type’ (*F*_(3,8379)_=11.4, *p*<.001) and ‘Choice Type’ × ‘Previous Feedback’ interaction (*F*_(3,8403)_=5.31, *p*<.001).

##### Learning Effect

Planned comparison using the Tukey HSD post-hoc test within ‘Learning’ factor split by ‘Choice Type’ revealed that pupil size was significantly increased for LP choices in ‘after learning’ condition compared with ‘no learning’ condition (Tukey HSD, *p*=.04) (Figure 5a, left panel). On the contrary, pupil size was significantly decreased for HP choices in ‘after learning’ condition (Tukey HSD, *p*<.001). Thus, for pupil size we observed the same pattern of the learning effects as that for RT.

**Figure 5.**
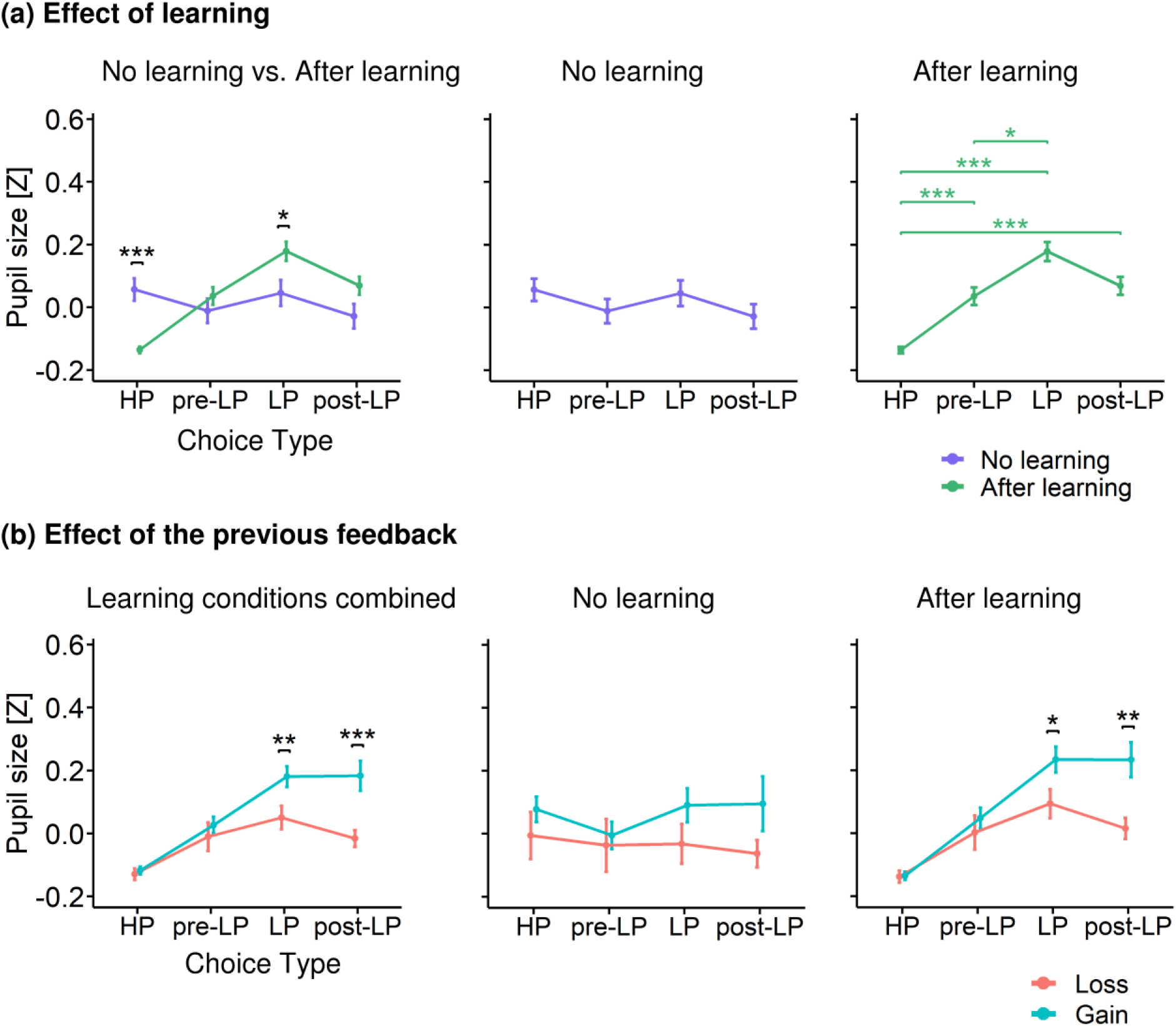
Pupil Size (z-scored) Represented as a Function of Choice Type. **(a)** Pupil size averaged within the interval −400–2200 ms relative to the response (button press) for different choice types under ‘after learning’ (green) and ‘no learning’ (slate blue) conditions. Left panel – pupil size differences between learning conditions within each choice type, middle and right panels – pupil size differences between choice types within ‘no learning’ and ‘after learning’ respectively. **(b)** Pupil size differences between previous outcomes within each choice type: ‘losses’ (salmon) vs. ‘gains’ (turquoise). Left panel – both learning conditions pooled together, middle and right panels – ‘no learning’ and ‘after learning’ conditions respectively. All other designations as in Figure 3.

Then we probed how ‘Choice Type’ influenced pupil size within ‘no learning’ and ‘after learning’ conditions analyzed separately. In contrast to ‘after learning’, in ‘no learning’ condition, there were no significant differences between choice types (Figure 5a, middle panel). Thus, in the absence of the internal utility model, pupil size did not depend upon the type of choice made by participants.

In ‘after learning’ condition (Figure 5a, right panel), pupil size was significantly greater during LP choices compared with HP choices (Tukey HSD, *p*<.001) and pre-LP choices (Tukey HSD, *p*=.011). Additionally, pupil size was significantly greater during pre-LP and post-LP choices compared with HP choices (Tukey HSD, *p*<.001). Thus, in convergence with the RT data, pupil size after learning was increased during disadvantageous choices compared to advantageous choices; it was also moderately yet significantly increased during advantageous choices on adjacent trials that immediately preceded and immediately followed disadvantageous choices (pre-LP and post-LP).

At least some of the disadvantageous choices were committed after losses and could be immediately caused by them. In order to exclude the possibility that this potentially retroactive mechanism could influence the results, we repeated the same analysis within the ‘Choice Type’ factor using a smaller restricted subset of data with uninterrupted sequences of pre-LP → LP → post-LP trials involving only gains on pre-LP trials. Comparisons in this reduced dataset closely reproduced the patterns of effects described above for a full dataset (supplementary materials, Figure S2). Thus, the effect of pupil dilation during disadvantageous choices was not caused by losses on trials preceding them.

In summary, we found that after learning, which led to formation of the utility model, two major changes occurred. First, pupil size became smaller for the repetitive HP choices, during which participants exhibited a stable preference for the advantageous stimulus, i.e., were keeping with a relatively safe strategy. Second, pupil size was increased for unsafe disadvantageous choices.

##### Effect of previous feedback

As with RT, the sign of the previous feedback differently affected pupil size depending on the Choice Type, as reflected by significant interaction ‘Choice Type’ × ‘Previous Feedback’, but the pattern was qualitatively different (Figure 5b, left panel). In contrast to RT, pupil size was greater after gains than after losses, but only during LP and post-LP choices (Tukey HSD, *p*=.023 and *p*=.002, respectively); both choice types implied a switch from one strategy to another (from exploration to exploitation in post-LP choice and from exploitation to exploration in LP choice). No significant differences in pupil size between gains and losses were observed for HP and pre-LP choices.

Again, to analyze ‘Previous Feedback’ influence on pupil size within ‘no learning’ and ‘after learning’ conditions separately, for illustrative purposes, we ran a similar LMM on ‘no learning’ and ‘after learning’ data subsets separately (Figure 5b, middle and right panels). The analysis revealed ‘Choice Type’ × ‘Previous Feedback’ significant interaction within ‘after learning’ only (*F*_(3,6823)_=3.94, *p*=.008): pupil size was greater after gains compared with losses for LP and post-LP choices (Tukey HSD, *p*=.023 and *p*=.002, respectively), but not in HP and pre-LP choices.

#### Subintervals within the Choice-Related Time Period

In the main analyses of the pupil size, we used a rather long choice-related time interval, which may be functionally heterogeneous. Therefore, we divided this time interval into three successive functional subintervals: prior to a choice (decision making and action initiation), between a choice response and feedback signal (internal outcome evaluation in anticipation of the feedback) and after feedback (matching expected and actual outcome). Then, we analyzed each subinterval independently using the same statistical procedure as that used for the full choice-related time interval (supplementary materials, Figure S3 and S4). In all three subintervals, we observed the pattern of results highly compatible with that obtained in the main analysis, with the latest subinterval (1000 – 2200 ms relative to the response) manifesting the most pronounced statistical effects. Importantly, in each of these subintervals, pupil size was greatest during disadvantageous trials, with a similar yet attenuated effect for advantageous choices on adjacent trials immediately preceding and immediately following disadvantageous trials – compared with the stable preference for advantageous choices (supplementary materials, Figure S3). Interaction ‘Choice Type’ x ‘Previous Feedback’ was significant in the second and third subintervals. Again, the pattern of results on these subintervals was similar to that obtained on a full response-related time interval of interest (supplementary materials, Figure S4). This suggested that a relatively increased pupil size reflected a protracted common process that affected different stages of decision making in regard to LP choices made during ‘after learning’ condition.

#### Pretrial Pupil Size

In order to evaluate whether the learning-induced effects found for choice-related pupil modulations involved slow tonic components and/or carry-over effects from previous trials, we additionally analyzed the pretrial time interval (−300 – 0 ms relative to fixation cross onset) using the same statistical procedure as that used for choice-related interval (Figure 6a). The following effects were statistically significant: ‘Choice Type’ (*F*_(3,7539)_= 10.98, *p*<.001), ‘Learning’ (*F*_(1,4504)_= 6.84, *p*<.001), ‘Learning’ × ‘Choice Type’ (*F*_(3,7535)_= 6.84, *p*<.001). Planned comparisons of ‘Learning’ × ‘Choice Type’ interaction revealed that learning success mainly increased pre-trial pupil size for post-LP choice (‘after learning’ vs ‘no learning’ for the post-LP: Tukey HSD, *p*<.001), but not for the LP choice itself (Figure 6a, left panel).

**Figure 6.**
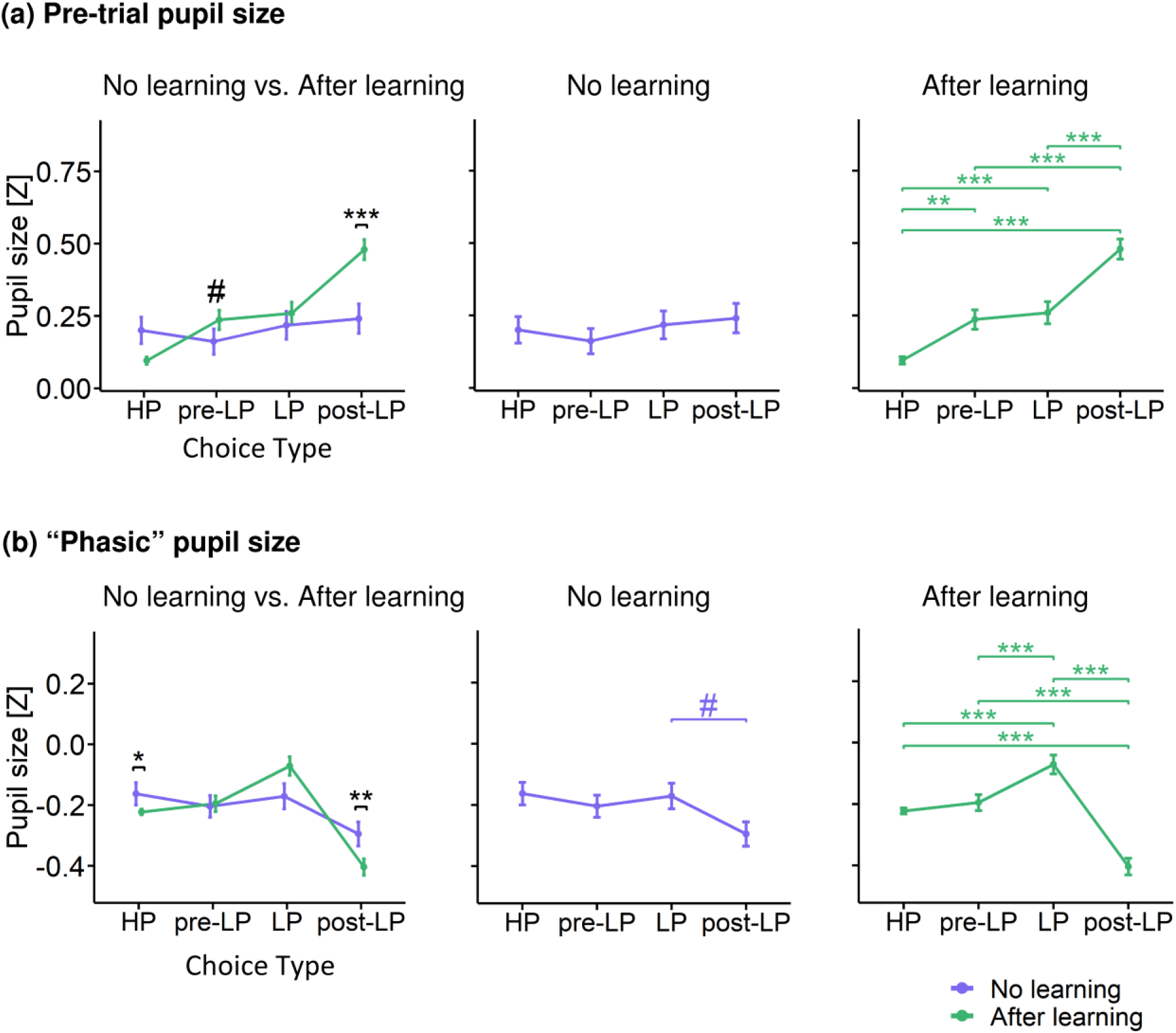
Baseline and Phasic Changes in Pupil Size (z-scored). **(a)** Pre-trial pupil size averaged within the interval −300–0 ms relative to the fixation cross onset for different choice types under ‘after learning’ (green) and ‘no learning’ (slate blue) conditions. **(b)** Phasic pupil size^†^ for different choice types under ‘after learning’ (green) and ‘no learning’ (slate blue) conditions. Left panel – pupil size differences between learning conditions within each choice type, middle and right panels – pupil size differences between choice types within ‘no learning’ and ‘after learning’ respectively. All other designations as in Figure 3. ^†^Note: Phasic pupil was calculated as the difference between the z-scored pupil size averaged within the interval −400–2200 ms relative to the response onset and pretrial pupil size. Pupil size was z-scored within each participant using all time points across all trials and pre-trial intervals. Consequently, subtraction between two values (pretrial and trial) produced negative z-scores for the phasic changes in the pupil size because of the pupillary light reflex evoked by luminance increment during the stimulus presentation.

In ‘no learning’ condition, pre-trial pupil size did not discriminate between choice types (Figure 6a, middle panel). In ‘after learning’ condition, the main choice-related distinction was a small “baseline” pupil size for HP choices as compared with pre-LP, LP and post-LP choices (Tukey HSD, *p*=.003, *p*<.001 and *p*<.001, respectively) (Figure 6a, right panel). Yet, the greatest pre-trial pupil size was observed not before the LP choice itself, but before post-LP choice (pre-LP VS LP after learning”: Tukey HSD, *p*<.001).

Thus, unlike pupil size during a choice, its modulations in the pre-trial period did not accentuate the impact of learning on pupil size during the disadvantageous choice. The dramatic increase in the “baseline” pupil size preceding post-LP choices is likely a carryover effect lasting from the choice-related pupil dilation in the previous LP trial (compare Figure 5a, right panel and Figure 6a, right panel). Yet, the fact that during ‘after learning’ condition the pupil was greater before pre-LP and LP trials compared with HP trials hints at some tonic effect, which contributed to disadvantageous choices.

Previous feedback did not affect pretrial pupil size in both learning conditions. Thus, the impact of previous feedback on the pupil size was mainly related to the choice response itself.

#### Baseline-Corrected Pupil Size

As an additional step, to corroborate the phasic nature of the pupil effects described in the main analysis, we used the conventional baseline-corrected choice-related pupillary response magnitude, and applied the same statistical model as in the main analysis. The following factors were significant: ‘Choice Type’ (*F*_(3,7529)_=16.87, *p*<.001), ‘Previous Feedback’ (*F*_(1,7524)_=8.69, *p*=.001), ‘Learning’ × ‘Choice Type’ interaction (*F*_(3,7522)_=3.97, *p*=.002).

Baseline correction preserved the basic pattern of results for the learning effect on choice-related pupil modulations except for the post-LP choices (Figure 6b). ‘Learning’ × ‘Choice Type’ interaction was partially due to the opposite direction of learning-induced pupil changes for the HP and LP choices, similar to that detected in the main analysis (Figure 6b, left panel). In addition, under ‘after learning’ condition, the maximal baseline-corrected pupil size was observed for the LP choices as compared to all the other choice types, which is also in concordance with the previous finding (compare Figure 6b, right panel with Figure 5a, right panel. Concurrently, baseline-corrected pupil size became minimal for post-LP choices - although apparently, this is a technical result of baseline subtraction. Inspection of Figure 6a, right panel shows that the pretrial pupil size for post-LP choices was enormously high, most likely due to the long-lasting impact of disadvantageous LP choice on the pupil size that sustained through the whole inter-trial period (compare Figure 6b, right panel with Figure 6a, right panel). The occurrence of profound technical interactions with the pre-trial baseline requires caution when using unsupervised usage of baseline correction for studying a “phasic” pupil response, especially when inter-trial interval is relatively short. To sum up, the additional analysis confirmed that the increased pupil size during disadvantageous choices made after learning involved a strong phasic component.

#### Correlational Analysis

Within ‘after learning’ condition, there was a significant negative correlation between the percentage of LP choices and LP-choice-related RT slowing (*r*_(73)_=-0.29, *p*=.01) (Figure 2b). In other words, the more often subjects made LP choices, the less RT slowed down in those choices compared with HP choices. Also, there was a similar correlation for pupil size (*r*_(73)_=-.25, *p*=.028) within the 1000-2200 ms time interval after response onset (Figure 2c): pupil dilation during LP choices was diminished in those participants who committed LP choices more often.

## Discussion

When offered a choice between two alternatives in a standard probability learning task, people occasionally shift their preference towards the option yielding a lesser (mathematical) expectation of the reward. To explain this suboptimal behavior, it has been hypothesized that people intentionally seek patterns in the sequences of outcomes, even though none are present and the probabilities of reward are held constant (see e.g., Ellerby & Tunney, 2017; Unturbe & Corominas, 2007). From this perspective, rare transitions from objectively advantageous to disadvantageous choices may represent a directed exploration strategy that guides the choice towards an option with uncertain payoff (Wilson et al., 2021).

To test this hypothesis, we contrasted pupil size and response time for LP and HP choices before and after the participants became aware of response-reward contingencies. We found that LP choices were linked to a profound RT slowing and greater pupil dilation, but only if the internal utility model has already been acquired by a participant. For the pupil size, this effect was strongly amplified for the LP choice, which immediately followed the gain as compared with loss in the preceding choice. Critically for the hypothesis tested, LP versus HP difference in pupil size involved a strong phasic component and was temporally related to the behavioral choice, but neither to the stimulus itself, nor to the feedback about the choice outcome.

Importantly, our analysis of probabilities of transitions from advantageous to disadvantageous choices proved that the disadvantageous choices themselves were not simply caused by negative outcomes of a preceding advantageous choice, i.e., most disadvantageous choices did not result from a simple Win-Stay Lose-Shift strategy (Ellerby & Tunney, 2017; Gaffan & Davies, 1981; Ivan et al., 2018).

In the following discussion, we will argue that the pattern of results obtained suggests that the rare, objectively disadvantageous choices, that violate the acquired internal utility model, do represent self-generated exploratory behavior. This exploratory strategy causes shifts in choice priorities in favor of information seeking, while its autonomic and behavioral concomitants are mainly driven by a conflict between the behavioral plan of the intended exploratory choice and its predominant alternative, which has already proven to be more rewarding during previous trials.

First, we wanted to ascertain that the observed RT and pupil size dynamics cannot be explained by general cognitive processes, non-specifically related to the value-based decision formation - sustained attention, internal error detection, outcome monitoring related to the external feedback, and reaction to a previous loss (for review see Zenon, 2019).

The simplest explanation of LP choices is the loss of attention to stimulus display during objectively disadvantageous choice. Response slowing was previously observed during continuous attentional tasks on the trials during which, according to participants’ reports, his/her attention was disengaged from the current task either due to the involvement in the inner thoughts or simply due to the decrement of alertness (Cohen & van Gaal, 2013; Dyson & Quinlan, 2003; O’Connell et al., 2009; Ratcliff & McKoon, 2008). However, in sharp contrast to the LP choices in our experiment, response slowing during unfocused attentional states was associated with reduced (not increased as during LP choices) task-evoked phasic pupil dilation (Figure 3a and 5a) (Unsworth & Robison, 2016). Convergent, instead of divergent, changes in the phasic pupil dilation and the response time during LP choices refute the suggestion that they originated from attentional lapses.

Still, another possibility is that response slowing and pupil dilation during LP choices are driven by internal detection of accidentally committed erroneous response. In the experimental tasks requiring participants to learn arbitrary association between visual stimuli and specific response, RT slowing accompanied with phasic pupil dilation is commonly observed not only after error commission (post-error slowing) (Critchley, Tang, Glaser, Butterworth, & Dolan, 2005; Wessel, Danielmeier, & Ullsperger, 2011), but also during the erroneous response itself (error slowing); such effect was tentatively ascribed to an error-evoked orienting response (Murphy, van Moort, & Nieuwenhuis, 2016). At first glance, “error-evoked” explanation seems plausible here, because relatively greater pupil dilation in the LP vs HP trials emerges as a result of a conscious appraisal of LP choices as disadvantageous (the effect was present exclusively in ‘after learning’ conditions, Figure 3 and 5), and hence “erroneous”. This “error-detection” explanation, however, is difficult to reconcile with our finding of highly significant pupil dilation and RT slowing in the prelude to an LP choice - a pre-LP trial (Figure 3 and 5), when no erroneous response was committed, and a participant undertook a “correct” advantageous choice. Also, in contrast to our data, pre-error speeding rather that slowing is commonly observed (Dudschig & Jentzsch, 2009), while we observed slower responses on pre-LP trials compared with HP trials.

The third cognitive process putatively involved in the LP choices could be external outcome monitoring; it implies that phasic pupil dilation is caused by the negative feedback contingent with the objectively disadvantageous choice (see e.g., Satterthwaite et al., 2007). Refuting this possibility, pupillary response during such choices emerges and sustains throughout almost the whole decision-making interval, long before the feedback was provided (Figure 4).

Apart from the cognitive processes involved in the LP choices themselves, a participant’s reaction to the negative outcome of the previous “correct” choice should also be considered as a possibility. Post-loss slowing has been reported for some gambling tasks (Brevers et al., 2015; Goudriaan, Oosterlaan, de Beurs, & van den Brink, 2005), although pupil measurements in these studies were lacking. In order to check whether the previous loss substantially influenced our data, we repeated the basic analysis on the RT and pupil size data using uninterrupted sequences of trials, in which the LP choices were committed exclusively after wins, i.e., after those HP choices that were rewarded (supplementary materials, Figure S1 and S2). In such a restricted dataset, the RT and pupil size effects remained highly significant, thus dismissing the predominant role of sensitivity to immediate previous loss in a subject’s decision to switch to the obviously disadvantageous choice.

After refuting alternative explanations of our RT and pupil findings, we argue that a participant’s decision to seek new alternatives (directed exploration) seems to be the most plausible explanation of behavioral and pupil changes evoked by spontaneous LP choices. This account is based on the findings in the literature that relate a slow RT and choice-evoked pupillary response to information processing and updating the internal model in the brain (see Zenon, 2019 for review). A critical distinction between the current and previous pupil studies of exploration/exploitation dilemma is the nature of exploratory choice itself. The previous pupillometric studies investigated so-called random exploration (Wilson et al., 2021): in these studies, exploratory choices were explicitly encouraged by a gradual decrease in the reward probability for a preferred choice in a restless multiarmed bandit task (Gilzenrat et al., 2010; Jepma & Nieuwenhuis, 2011). Our experimental design was fundamentally different, as it did not involve any systematic changes in the utility of a particular choice and the decider had known the probability of likely outcomes from the previous experience. Thus, our findings provide the first evidence for pupillary response accompanying self-generated exploratory decisions such that participants intentionally choose a risky exploratory option against their behavioral bias toward value-driven choices.

Although the differentiation between random (purely uncertainty-driven) and self-generated or directed exploration (intentional information seeking) is a long-standing problem in psychological literature (Berlyne, 1966; Wilson et al., 2021), physiological concomitants of the directed exploration are largely unknown (but see Zajkowski et al., 2017). In this respect, our findings add an important new dimension to the existent knowledge about the relation of pupil dilation as a measure of LC-NA arousal to human exploratory behavior. Moreover, inclusion of ‘no learning’ and ‘after learning’ conditions in the present study allowed us to examine the change in choice-related pupillary response from random to self-generated exploration.

The question we were seeking to answer was whether the concept of subjective uncertainty/surprise used to explain pupil-related LC-NA arousal during exploratory choices (Van Slooten et al., 2018) can also be applied to self-generated exploration, or whether different decision processes are invoked depending on the source of uncertainty.

One possibility is that the effect of uncertainty played a similar role in producing transient pupillary responses during both random and self-generated exploratory choices. Hypothetically, in ‘no learning’ condition, during which the reward structure remained largely unknown for participants, either choice was “random” and was characterized by an equal uncertainty in the prior belief regarding the outcome. This can explain why a pupillary response to either choice did not distinguish the LP and HP choices in ‘no learning’ conditions (Figure 5а, middle panel). As soon as the participants learned to prefer choices that had been probabilistically associated with positive outcomes (‘after learning’ condition), the uncertainty was greatly reduced for such advantageous choices, leading to a highly significant attenuation of both choice-related pupillary response and response time costs (Figure 3a and 5a). Still, ambiguity remained whether other response strategies incorporating occasional risks – choices with a low payoff probability – might produce better total outcomes than the status quo. On the basis of the empirical consensus of association between pupillary response and subjective uncertainty (Richer & Beatty, 1987; Satterthwaite et al., 2007; Urai et al., 2017; Van Slooten et al., 2018), one might predict that the choice-related increase in pupil size, although attenuated for safe choices after learning, would be preserved for the risky explorative ones. This learning-related difference in pupillary responses between the safe and risky choices was exactly what we observed in our data (Figure 5a).

Thus, at first glance, our findings match well with the principle derived from computational modelling (Jepma & Nieuwenhuis, 2011; Urai et al., 2017; Van Slooten et al., 2018) – phasic pupil dilation is proportional to the subjective estimate of uncertainty. On the other hand, in our experimental settings, the implicit conflict between “safe” and “risky” explorative options is an inevitable consequence of self-generated exploratory choice. The learned value of the objectively advantageous choice is known to create an unconscious value-driven bias (for review see Anderson, 2016) that can interfere with the effects of voluntary endogenous selection determined by the goal of a subject (Preciado, Munneke, & Theeuwes, 2017). Implicit conflict with this unconscious bias arises when a preferable “safe” action plan is overruled by a deliberate “risky” exploratory decision. Previously, the phasic increase in pupil size was found to be a robust measure of implicit conflict between task-appropriate and habitual automatic responses in a color-naming Stroop task (Laeng et al., 2011). In the context of a value-driven choice, pupil dilation was mainly studied under an explicit conflict, which was parametrically manipulated by changing the already learned differences in the likelihood of reward between two alternatives. Specifically, phasic pupil dilation was found to closely track a degree of explicit conflict, being inversely proportional to the difference in the reward probability for each of the alternative options (Van Slooten et al., 2018). A similar effect was described for the intertemporal choice paradigm when the degree of conflict between the competing subjective preferences for immediate or delayed reward was also parametrically manipulated and formally modelled (Lin et al., 2018). Notably, strongest dependency of pupillary response and decision time cost on the degree of explicit conflict was found for appetitive conditions, i.e., a choice between two conflicting equally desirable win-win action plans (Cavanagh et al., 2014). Thus, when each alternative has significant advantages and disadvantages, people often experience conflict that makes the choice aversive and causes choice-related pupil dilation.

A strong influence of explicit conflict between the two action plans on pupillary response and RT supports our hypothesis that implicit conflict pertinent to self-generated exploratory choices has a similar effect on pupil dilation. In the latter case, conflict arises from competition for action selection between the unconsciously biasing effect of previously rewarded action and voluntary decision, which is shifting choice priorities in favor of information seeking.

We therefore tried to distinguish between “subjective uncertainty” and “conflict” effects by analyzing the pupil dilation in the “safe” choices that immediately preceded and followed the deliberate “risky” choice, as well as by considering the effects of the previous feedback sign on pupillary response. First, both behavioral and pupil results suggest a protracted decision formation such that an intentional exploratory decision was actually made during the preceding trial, maintained, and then enacted during exploratory choice itself (Figure 3a and 5a). This finding is fully compatible with the previous reports demonstrating that pupillary response associated with the internal state of random exploration begin to develop on the trial preceding the exploratory choice itself (e.g., Gilzenrat et al., 2010; Jepma et al., 2010; Jepma & Nieuwenhuis, 2011). However, the pure uncertainty account is difficult to reconcile with our finding that phasic pupil response and response time remained relatively high on the trial immediately following self-generated exploratory choice, when a participant returned to the “safe” choice strategy with a knowingly high payoff probability (Figure 3a and 5a). Neither, effect of uncertainty alone can explain why self-generated exploratory choice (during ‘after learning’ condition) elicited greater pupil dilation and slower response time than a random exploratory choice with completely unpredictable choice outcome (during ‘no learning’ condition) (Figure 3a and 5a). This rather suggests a cumulative contribution of uncertainty regarding the desired outcome and a conflict with a value-driven bias in the self-generated exploration.

Further evidence for the contribution of conflict dimension to the self-generated exploratory choice is provided by the amplifying role of the previously obtained positive feedback on the increased pupillary response. The previous reward as compared to punishment was associated with a greater pupil dilation in both “risky” explorative choices and “safe” post-explorative choices, while this effect was absent for the two other “safe” advantageous choice types (HP-choice and pre-LP choice) (Figure 5b). Importantly, in both choice types sensitive to the previous reward, the participants shifted to a response that was incongruent with the positive outcome of the previous choice. They changed their action plan towards exploration after being rewarded for the exploitative action (LP-choice), or, vice versa, returned to exploitative strategy despite the successful outcome on the previous, explorative, action (post-LP choices). Since the other “safe” choices followed uninterrupted history of the previous frequently rewarded choices, they did not conflict with a previously rewarded action (supplementary materials, Figure S2). This finding indicates that the pupil dilation during self-initiated exploration is likely to reflect more than one process occurring concomitantly.

Notably, for RT measurements, the effect of the previous reward during explorative choices and post-explorative choices was not observed (Figure 3b). This may be explained by the powerful effect of the preceding loss on response time – i.e., post-loss slowing (Brevers et al., 2015; Goudriaan et al., 2005) – but not on pupil size (Figure 5b) that was seen in both the HP- and pre-LP choices. This generally adaptive tendency, which serves to promote caution in decision making after losses, may counteract the opposite effect of previous reward on response speed in LP and post-LP trials.

To sum up, while subjective uncertainty is likely to play an important role in phasic pupil dilation caused by directed exploration, it is hardly the only factor determining strong enhancement of pupil size observed here in such choices. Pupil dilation is rather caused by the combined effect of subjective uncertainty regarding exploratory choice outcome and the conflict signal broadcasting that the intended exploratory action violates the internal utility model, which favors the frequently rewarded alternative.

The self-generated decisional challenge whether to explore a set of alternative choices or stick to the opportunity to make a ‘default’ choice suggests a comparison process taking place within the anterior cingulate cortex (ACC). Given that ACC in generally involved in the comparison between the outcome values of different choice options (Kolling, Behrens, Wittmann, & Rushworth, 2016), and specifically in conflict monitoring processes (Botvinick, Cohen, & Carter, 2004; Shenhav, Botvinick, & Cohen, 2013), the detection of conflict with the inner utility model in self-initiated exploratory choices may drive transient changes in LC-NA-mediated arousal, which, in turn, increases phasic pupil dilation observed here.

This speculation is consistent with the recent monkey study directly demonstrating that in some cases, pupil related modulations of spontaneous neuronal activity reflect signals occurring first in ACC and then being transmitted to the LC and other subcortical and cortical structures (Joshi, Li, Kalwani, & Gold, 2016). It is also in accord with the human findings showing that the phasic pupil dilation during explicit high-conflict appetitive choices correlates with increased mediofrontal activation (Cavanagh et al., 2011; Cavanagh et al., 2014). Therefore, phasic pupil dilation triggered by the implicit conflict in our experiment may reflect downstream signal of conflict processing in ACC.

Evidence from multiple psychological and physiological research indicate that conflict is emotive, and triggers a negatively valenced affective state accompanied by changes in heart rate, skin conductance, body temperature, pupil response, and muscle tone (for review see Saunders, Lin, Milyavskaya, & Inzlicht, 2017). One cannot exclude, therefore, that the negative emotion triggered by implicit conflict may contribute to pupil dilation we observed for self-generated exploratory choices. By changing the inner affective state, learning from the history of rewards and punishments may reach perception of knowing without conscious awareness (Bechara, Damasio, Tranel, & Damasio, 1997) – the process of nonconscious information processing that has been referred to as System 1 (Kahneman, 2003). Functionally, changes in the inner physiological state may serve as a subconscious warning signal informing a decision maker that his/her deliberate action plan violates the inner brain model for utility of the intended action.

## Conclusion

The study demonstrates that response slowing and augmented pupil-related phasic arousal characterize a participant’s decision to take risk of information seeking by choosing an uncertain alternative over the rewarding one in a two-choice probabilistic learning task. The behavioral and pupillometric findings also suggest that such directed exploration is bound to a conflict between the deliberate explorative choice with uncertain outcome and the inner bias to select the option with the highest value. The close relationship between self-generated choices and pupil-related arousal makes a simple probabilistic learning task a complementary instrument for studying neural underpinnings of directed exploration and the underlying pathophysiology of its abnormalities in mental disorders.

## Acknowledgments

We are grateful to Ivan S. Pozdniakov for invaluable help in recruiting participants.

## Availability of data and materials (Open Practices Statement)

Data are openly available at https://doi.org/10.6084/m9.figshare.16825570. The experiment was not preregistered.

## Ethics approval

The study was conducted following the ethical principles regarding human experimentation (Helsinki Declaration) and approved by the Ethics Committee of the Moscow State University of Psychology and Education.

## Consent to participate

All participants signed the informed consent before the experiment.

## Authors’ contributions

GK, AP, TS, BC contributed to conception and design of the study. GK, KS, AP, AR were responsible for the experiment and data acquisition. GK, KS, AP, VM performed statistical analysis of the data. KS, VM, AR, TS, BC prepared illustrations. GK, KS, VM, TS, BC interpreted the data. GK, BC wrote the first draft of the manuscript. GK, KS, VM, AR, TS, BC wrote sections of the manuscript. All authors contributed to manuscript revision, read and approved the submitted version.

