## Supplementary materials for "Pupil Dilation and Response Slowing Distinguish Deliberate Explorative Choices in the Probabilistic Learning Task"

### Analysis of pupil size in uninterrupted sequences of pre-LP → LP → post-LP trials involving only gains on pre-LP trials

In order to check whether the effect of RT slowing during disadvantageous LP choices was not a consequence of negative outcomes of a preceding advantageous choice, we repeated the main analyses using a smaller restricted subset of data within uninterrupted sequences of pre-LP → LP → post-LP trials involving only gains on pre-LP trials. All basic findings concerning the effects of ‘Learning’ and ‘Choice Type’ were reproduced both for the RT (Figure S1) and the pupil size (Figure S2). Thus, the effect of pupil dilation during disadvantageous choices was not caused by losses on trials preceding them.

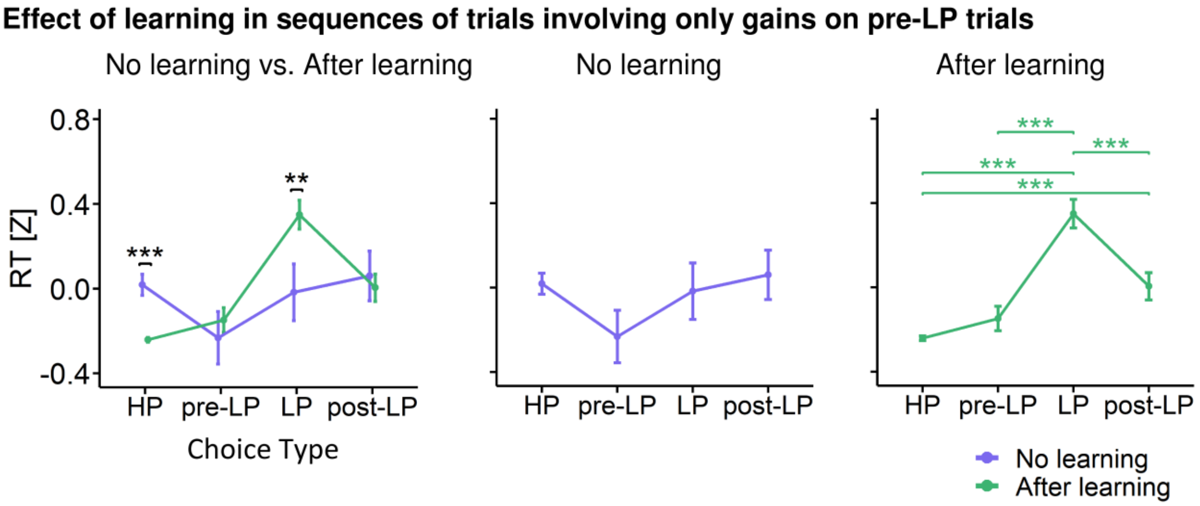


**Figure S1** Response time in dependence on learning condition as a function of choice type in uninterrupted sequences of pre-LP → LP → post-LP trials involving only gains on pre-LP trials. All designations as in Figure 3 in the main text.

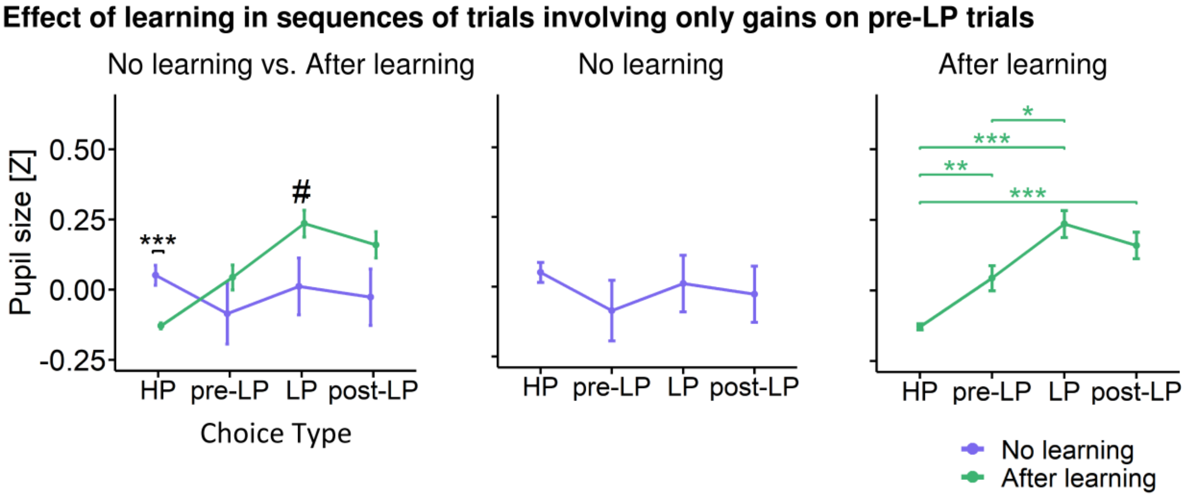


**Figure S2** Pupil size (z-scored, averaged within the interval -400–2200 ms relative to the response) in dependence on learning condition as a function of choice type in uninterrupted sequences of pre-LP → LP → post-LP trials involving only gains on pre-LP trials. All designations as in Figure 3 in the main text.

**Analysis of pupil size using subintervals within the time interval of interest**

In the main analyses of the pupil size, we used a rather long response-related time interval, which may be functionally heterogeneous. Therefore, we divided the above time interval into three functional subintervals (Figure S3 and S4).

For the first subinterval related to decision making and action initiation (-400–0 ms relative to response) factor ‘Previous Feedback’ ($F_{(1,8367)}$=6.79, $p$=0.009) and the ‘Choice Type’ × ‘Learning’ interaction (($F_{(3,8367)}$=6.79, $p$<0.001) were significant. The difference between ‘after learning’ and ‘no learning’ was present only within HP choices: pupil size was greater within ‘no learning’ condition (Tukey HSD, $p$<0.001) (Figure S3a). Pupil was dilated in pre-LP ($p$=0.003), LP ($p$<0.001) and post-LP choices ($p$<0.001) compared with HP choice, and such differences were observed ‘after learning’ only (Figure S3a). Pupil size was more dilated after gains than after losses relatively equally within all choice types (Figure S4a).

For the second subinterval related to internal outcome evaluation and feedback anticipation (0–1000 ms) significant factors were 'Choice Type' ($F_{(3,8376)}$=4.9, $p$=0.0018), ‘Previous Feedback’ ($F_{(1,8390)}$=4.9, $p$=0.002), their interaction ($F_{(3,8371)}$=4.38, $p$=0.0016), ‘Choice Type’ × ‘Learning’ interaction (($F_{(3,8362)}$=8.98, $p$<0.001). Post hoc comparisons revealed the same pattern as in the previous interval (Figure S3b). In addition, pupil size after gains was greater than after losses during LP (Tukey HSD, $p$=0.007) and post LP choices ($p$<0.001) (Figure S4b).

For the third subinterval related to matching expected and actual feedback (1000–2200 ms), the same factors and their interactions were significant: 'Choice Type' ($F_{(3,8384)}$=12.3, $p$<0.001), ‘Previous Feedback’ ($F_{(1,8413)}$=4.9, $p$<0.001), their interaction ($F_{(3,8387)}$=7.24, $p$<0.001), ‘Choice Type’ × ‘Learning’ interaction (($F_{(3,8370)}$=12.28, $p$<0.001). Within this subinterval the difference between ‘no learning’ and ‘after learning’ conditions was observed not only within HP choice ($p$<0.001), but also within LP choice ($p$=0.0086) (Figure S3c). Within the ‘after learning’ condition pupil size differentiated HP choice from all other choices, and was greater during LP choice than during pre-LP choice. ‘Previous Feedback’ results reproduced: pupil was dilated after gains compared with losses in LP (Tukey HSD, $p<0.001$) and post-LP choices ($p<0.001$) (Figure S4c).

Although this was not the aim of the current study, we also evaluated the influence of the ‘Current Feedback’ factor on the pupil size during the third subinterval related to matching expected and actual feedback. No significant effects were found in both ‘No learning’ and ‘After learning’ conditions for the ‘Current Feedback’ factor ($F_{(1,1509)}= 0.19$, $p=0.65$ and $F_{(1,6797)}= 1.55$, $p=0.21$, respectively), as well as for ‘Choice Type’ × ‘Current Feedback’ interaction ($F_{(3,1507)}= 0.72$, $p=0.53$ and $F_{(3,6808)}= 0.29$, $p=0.83$, respectively).

Thus, in all the three subintervals, post hoc testing within the 'Choice Type' factor produced the pattern of results highly compatible with that obtained on the full response-related time interval of -400 – 2200 ms, with the third subinterval manifesting the most pronounced statistical effects (Figure S3c). Importantly, in all of these subintervals, pupil size was greatest during the LP choices. Interaction 'Choice Type' x ‘Previous Feedback’ was significant in the second and third subintervals. Again, the pattern of results on these subintervals was similar to that obtained on a full response-related time interval of interest (Figure S4b and S4c).

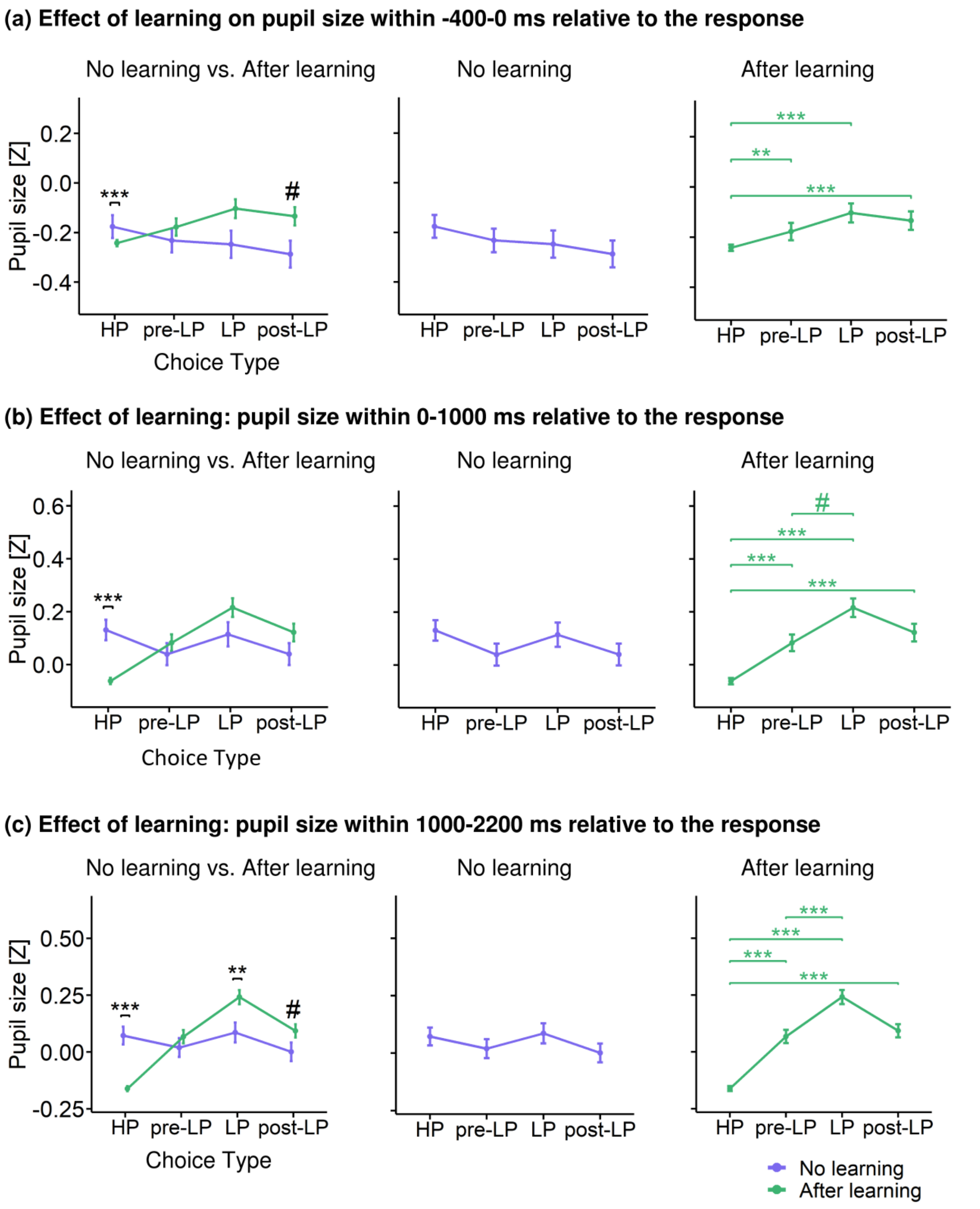


**Figure S3** Pupil size modulations by the learning condition as a function of choice type, with an average pupil size estimated within the subintervals **(a)** -400–0 ms, **(b)** 0–1000 ms, and **(c)**

1000–2200 ms relative to the response (button press). All designations as in Figure 3 in the main text.

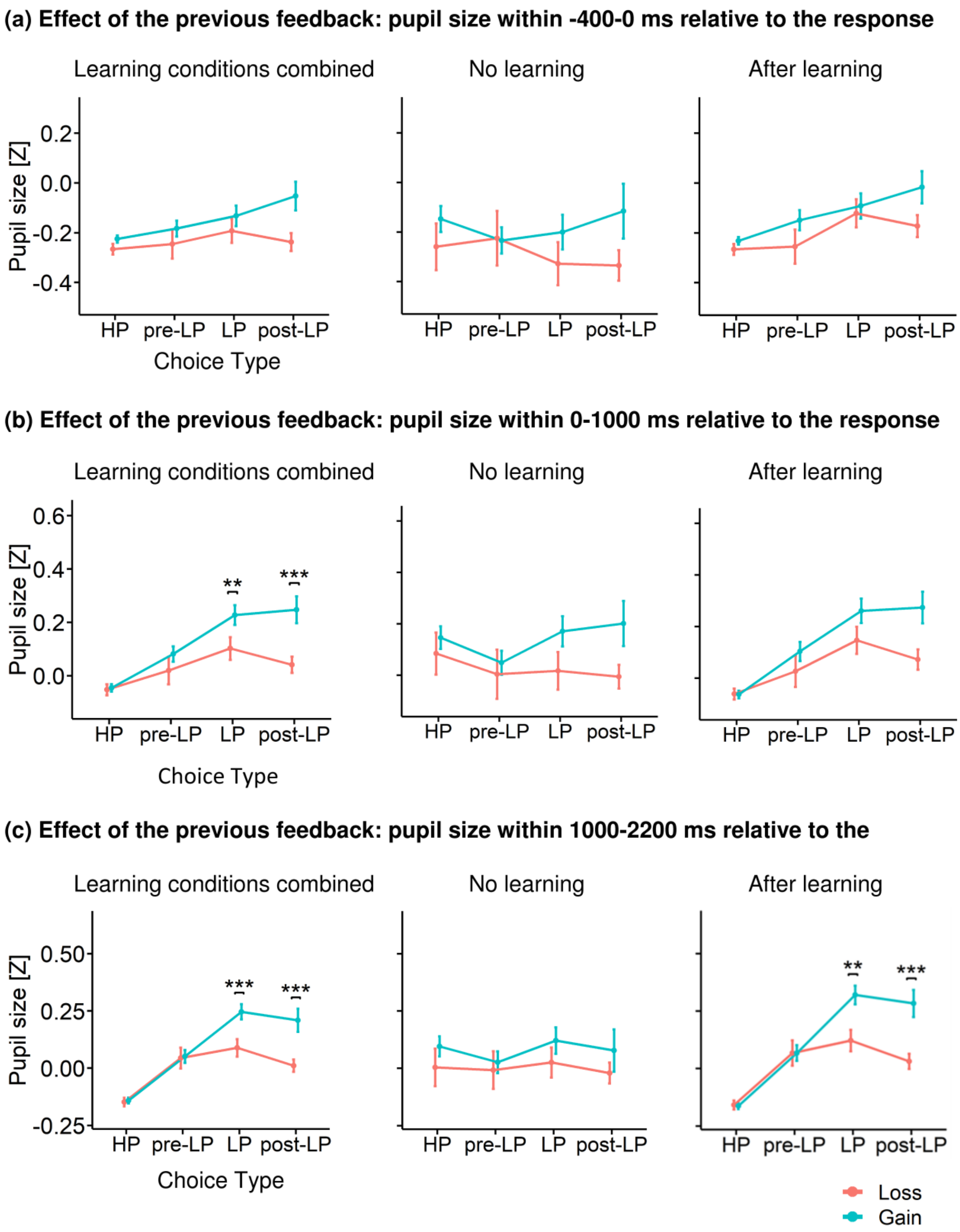


**Figure S4** Pupil size modulations by the sign of the previous feedback as a function of choice type, with an average pupil size estimated within the subintervals **(a)** -400–0 ms, **(b)** 0–1000 ms, and **(c)** 1000–2200 ms relative to the response (button press). All designations as in Figure 3 in the main text.
